# Molecular and cellular determinants of motor asymmetry in zebrafish

**DOI:** 10.1101/666594

**Authors:** Eric J. Horstick, Yared Bayleyen, Harold A. Burgess

## Abstract

Asymmetries in motor behavior, such as human hand preference, are observed throughout bilateria. However, neural substrates and developmental signaling pathways that impose underlying functional lateralization on a broadly symmetric nervous system are unknown. Here we report that in the absence of over-riding visual information, zebrafish larvae show intrinsic lateralized motor behavior that is mediated by a cluster of 60 posterior tuberculum (PT) neurons in the forebrain. PT neurons impose motor bias via a projection through the epithalamic commissure to the habenula. Acquisition of left/right identity is disrupted by heterozygous mutations in *mosaic eyes* and *mindbomb*, genes that regulate Notch signaling. These results define the neuronal substrate for motor asymmetry in a vertebrate and support the idea that developmental pathways that establish visceral asymmetries also govern acquisition of left/right identity.

## Introduction

In many bilaterian organisms the nervous system shows striking structural and functional lateralization, and many species show prominent asymmetric motor behaviors such as hand or paw preference (Rogers, 2009). However, attempts to link neuroanatomic and motor asymmetries have yielded contradictory and inconclusive results in humans (Good et al., 2001; Guadalupe et al., 2014; Sun et al., 2012). In other species, despite correlations between asymmetric properties of the nervous system and lateralized behavior, causal relationships have not been established (Davison et al., 2009; Gutierrez-Ibanez et al., 2011; Jozet-Alves et al., 2012; Lee et al., 2017). In the absence of a compelling neuronal basis, it has been difficult to resolve molecular determinants that drive the development of lateralized behavior.

Research to date has produced limited insight into molecular mechanisms that establish motor asymmetries. Consistent with early theories that outlined a role for single genes of large effect, twin studies on the heritability of handedness in humans revealed an important genetic component (Annett, 1972; McManus, 1985; Medland et al., 2006). However, genome-wide genetic studies failed to uncover loci with large contributions (Armour et al., 2014; Eriksson et al., 2010; Somers et al., 2015). Behavioral hand-preference is established as early as 10 weeks of gestation in humans and left/right asymmetries in gene expression within the spinal cord have been identified at this stage, suggesting a role for genes that pattern the nervous system (Hepper et al., 1998; Ocklenburg et al., 2017). Abnormalities in neuroanatomical asymmetry and handedness are associated with schizophrenia and other neurodevelopmental disorders, however without knowledge of the underlying neuroanatomical substrates or developmental pathways these findings are difficult to interpret (Dean et al., 2016; Markou et al., 2017; Sommer et al., 2001).

In the absence of a clear structural basis for a vertebrate motor asymmetry, many studies have instead focused on molecular genetic pathways that govern the development of brain asymmetries. In particular, work in zebrafish has outlined molecular genetic pathways that govern asymmetric morphogenesis and gene expression within the dorsal diencephalon. In zebrafish, as in many other vertebrates, the habenula shows a pronounced hemispheric asymmetry with well characterized differences in the size, composition and connectivity of subnuclei on the left and right sides (Aizawa et al., 2007, 2005; Amo et al., 2010; Concha et al., 2000; Gamse et al., 2005). Moreover, functional differences between hemispheres are also apparent, with olfactory cues preferentially activating the right habenula and visual stimuli driving responses in the left habenula (Cheng et al., 2017; Dreosti et al., 2014; Krishnan et al., 2014; Zhang et al., 2017). Behaviorally, the habenula in fish has been implicated in social conflict, anxiety and avoidance-learning (Amo et al., 2014; Chou et al., 2016; Facchin et al., 2015; Lupton et al., 2017) and the left-habenula in specifically in attenuating fear and driving light-preference behavior (Duboue et al., 2017). However, to date, assays reporting individual lateralized motor behavior in zebrafish have proven to difficult to reproduce, precluding efforts to resolve underlying asymmetries in brain structure (Barth et al., 2005; Facchin et al., 2009).

Here we report that individual larval zebrafish show a consistent motor asymmetry across multiple behavioral assays when tested in the absence of visual stimuli. Motor identity is maintained by a cluster of 60 neurons in the rostral lobe of the posterior tuberculum that project to the habenula nuclei, ablation of which also degrades motor bias. Finally, we demonstrate that lateralized behavior is disrupted by haploinsufficient mutations in genes that regulate Notch signaling, supporting the idea that the same pathway that establishes visceral asymmetries during development also governs acquisition of individual left/right motor identity.

## Results

### Larval zebrafish show persistent individual lateralized behavior in locomotor trajectories

After loss of illumination, 6 days post-fertilization (dpf) larval zebrafish initiate a circular swimming behavior in which they repeatedly perform same-direction turn maneuvers in a restricted spatial area (Figure 1A)(Horstick et al., 2017). Same-direction turn movements were sustained in individual larvae for two minutes (Figure 1B), however across the population there was no net tendency for larvae to preferentially circle to the left or right (mean Net Turn Angle (NTA) = −60.8 ± 84.2, one-sample t-test against 0, p=0.24; Figure S1A). However, we asked whether individual larvae show lateralized behavior — preferentially swimming in a left or right direction — or randomly select a circling direction on each event. We therefore probed individuals with a series of four light-off trials, each separated by several minutes of illumination. Larvae that circled in a rightward direction on the first trial, showed a significant tendency to circle rightward on subsequent trials, and similarly, left-circling larvae continued to show a leftward bias (repeated measures ANOVA, main effect of trial 1 direction F_(1,65)_=22.20, p<0.001; Figure 1C). Lateralized behavior was not observed when larvae were tested under constant illumination, as groups initially classified as left/right based on the first trial did not show differences in mean direction on subsequent trials (F_(1,61)_=1.08, p=0.30; Figure 1D). Directional bias on dark trials manifest as a 69.6% probability for larvae to circle in the same direction as on the first trial (match index, p < 0.0004 compared to 50% probability; Figure 1E). Circular swimming is primarily driven by routine-turn maneuvers (Horstick et al., 2017). We therefore used high-speed video recordings and kinematic analysis to directly measure routine-turn direction across a series of four light off trials (Figure 1F). Again, we noted that turn direction on trials 2-4 was significantly correlated with turn direction on the first trial (Figure S1B). Next, we calculated the percentage of routine-turns made in a rightward direction on each trial, then used the mean of the four trials to represent each larva’s direction preference. In the dark, the distribution of direction preferences strongly deviated from the expected distribution: 41% of larvae (37/89) produced fewer than than 24% or greater than 76% rightward turns, whereas if turn-direction was random on each trial, only 10% of larvae would have shown this level of bias (Monte Carlo simulations, p < 0.0001; Figure 1G). In contrast, under constant illumination the distribution of routine-turn direction bias was similar to the expected distribution (p=0.119; Figure 1H). These results reveal that zebrafish larvae raised in the same environment stochastically acquire a left/right directional bias in motor behavior that is manifest when tested under dark conditions.

**Figure 1:**
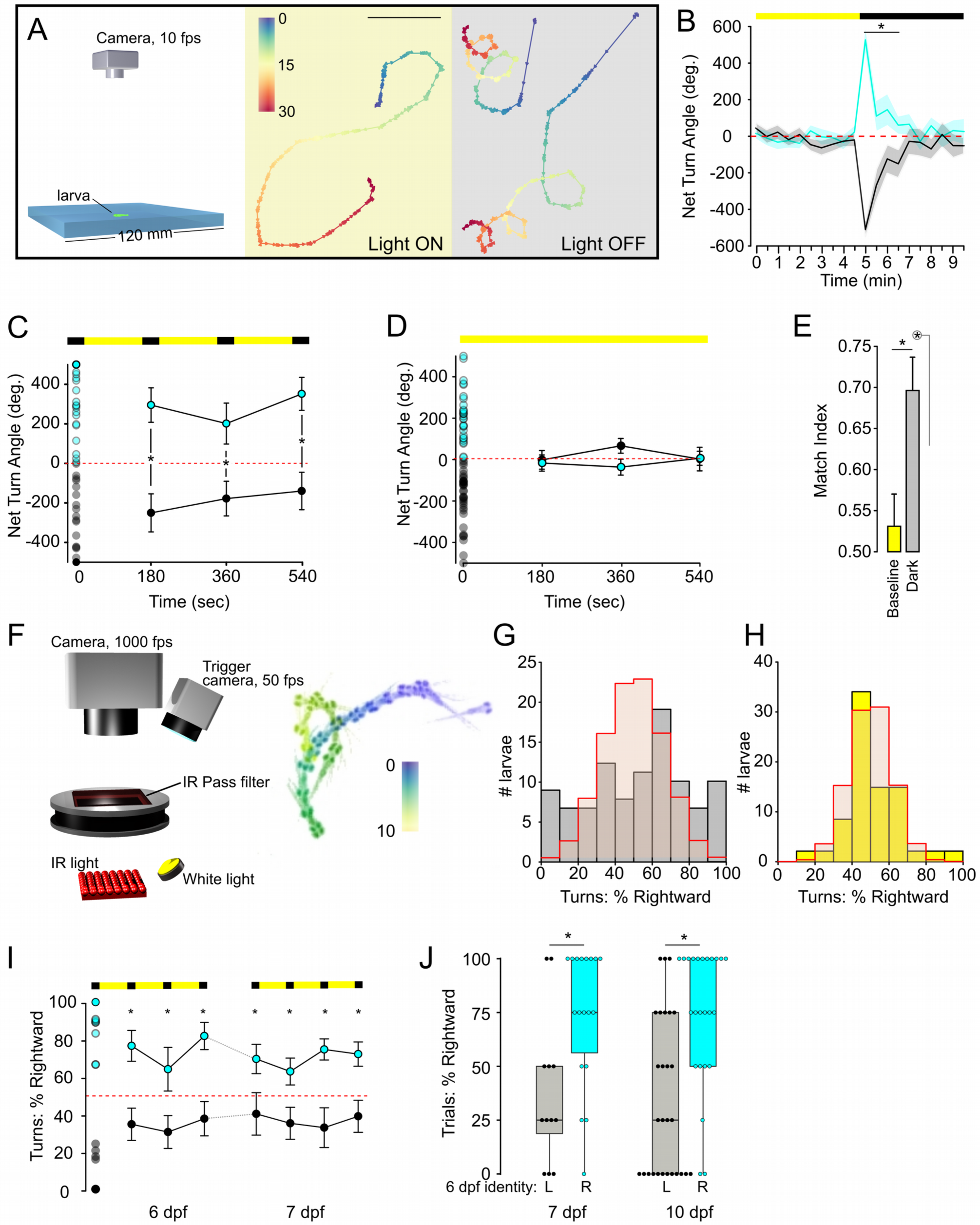
Individual larvae show persistent motor asymmetry during dark-induced circling behavior. A. Schematic of setup for path trajectory analysis (left). Swim trajectories for single larvae during baseline illumination and immediately after the loss of light (yellow and grey backgrounds respectively). Color scale represents time (seconds). Arrowheads along path line represent orientation of larva. Scale bar 20 mm. B. Net turn angle over 30 s intervals for larvae before and after the loss of illumination (indicated by top horizontal bars). Individuals were classified as either right (cyan, N=25) or left biased (grey, N=34) based on the first 30 s interval after the loss of illumination. * p < 0.05, t-test between groups. C. Net Turn Angle for larvae over four 30 s light off trials. Horizontal bars at top indicate baseline illumination (yellow) and light-off (black) periods. Open circles in the first trial (time 0) represent individual NTA for all larvae tested and were used to classify larvae as right (cyan, N=34) or left-biased (black, N=34). Subsequent points represent mean and standard deviation for left/right groups on trials 2-4. * p < 0.05, t-test between groups. D. As for (C) during constant illumination (right, N=29; left, N=35). E. Match Index during baseline illumination (yellow, N=64) and after the loss of illumination (grey, N=68). * p < 0.05, Mann-Whitney U test and (circled) 1-sample permutation test to 0.5. F. Visually isolated chamber for high-speed recordings that enable kinematic analysis of routine turns during circling behavior. Right: Time lapse montage over 10 s interval following loss of illumination. Color indicates time. G-H. Histogram for percentage of turns executed in a rightward direction (mean of four 10 s trials) after loss of illumination (G, grey, N=89 larvae) or during constant illumination (H, yellow, N=39). Red line represents expected distribution from Monte Carlo simulation assuming larvae have no directional bias. I. Rightward turn preference over 24 hour interval. Larvae were classified as right (cyan, N=14) or left (black, N=12) biased by four repeated dark recordings at 6 dpf. In this analysis, larvae with less than 33% of turns in a rightward direction were classified as left-biased, and those with greater than 66% classified as right-biased. At 7 dpf % rightward turn use was measured in L/R classified groups. Dotted red lines at 50% indicates no bias.* p < 0.05 between groups. J. Percentage of trials that had a net rightward bias for larvae tested at 7 or 10 dpf (N=30, 52 respectively), after being classified as left or right-biased at 6 dpf. * p < 0.05, Mann-Whitney U test.

A form of unstable lateralized eye-use behavior has been reported in zebrafish, with individuals switching eye-preference over several minutes (Andrew et al., 2009). In contrast we found that turn-bias during dark-induced circling behavior was sustained for at least 45 min (Figure S1C). Next, we tested individuals at 6 dpf to establish their left/right motor identity, then returned them to an incubator overnight. The next day, we recorded responses to a second set of four light-off stimuli. Turn direction bias for individual larvae was significantly correlated between days (Spearman *rho*=0.58, p=0.0009; Figure 1I-J, S1D) and also sustained in larvae that were raised for 4 days between tests (*rho*=0.39, p=0.0003; Figure 1J, S1F). Decomposing turn bias into its magnitude and direction components revealed that overall left/right direction preference was maintained (d6 to d7, Mann-Whitney U p=0.012; d6 to d10, p=0.00039; Figure 1J) although the strength of the preference was not correlated across days (d6 to d7, *rho*=-0.02, p=0.9; d6 to d10, *rho*=0.054, p=0.7; Figure S1E,G). Thus left/right motor identity is sustained in individual larvae for several days. In birds stable perceptual asymmetries are conferred by visual experience during embryonic development (Rogers, 1982). In contrast, dark-reared zebrafish larvae showed normal lateralized behavior, excluding an instructive role for visual experience in the acquisition of motor identity (Figure S1H). Together, these data provide compelling evidence that zebrafish larvae show a robust motor asymmetry, manifest as persistent individual differences in the direction of circling behavior after loss of illumination. Behavioral asymmetry in zebrafish is of moderate strength: larvae initiate movement in their preferred direction on around 70% of trials. Similar numbers of larvae are left and right-biased, with no apparent population-level asymmetry.

Next we asked whether motor bias is also present in responses to other stimuli. Zebrafish show positive phototaxis, and select and swim toward one of two simultaneously illuminated regions (Burgess et al., 2010). We first classified larvae as left or right biased during dark-induced circular swimming, then used a closed-loop system to present freely swimming larvae with two identical target spots on the left and right (Figure 2A,B). Target choice was positively correlated with circular swimming direction (Figure 2C). We also tested whether lateralized motor behavior occurred during responses to a non-visual stimulus. In zebrafish, intense auditory stimuli elicit escape responses that are initiated with a rapid bend to either the left or right, raising the possibility that startle direction might show a motor asymmetry. We again pre-classified larvae using dark-induced circular swimming, then measured startle direction. Under constant illumination, escape response direction showed little or no correlation with circular swim direction (Figure 2D, yellow background). However auditory stimuli that were presented in the dark evoked startle responses whose direction showed a significant match to the direction of circular swimming (Figure 2D, grey background). Thus, motor bias was apparent in three different assays, consistent with individual larvae having an intrinsic left/right motor identity.

**Figure 2:**
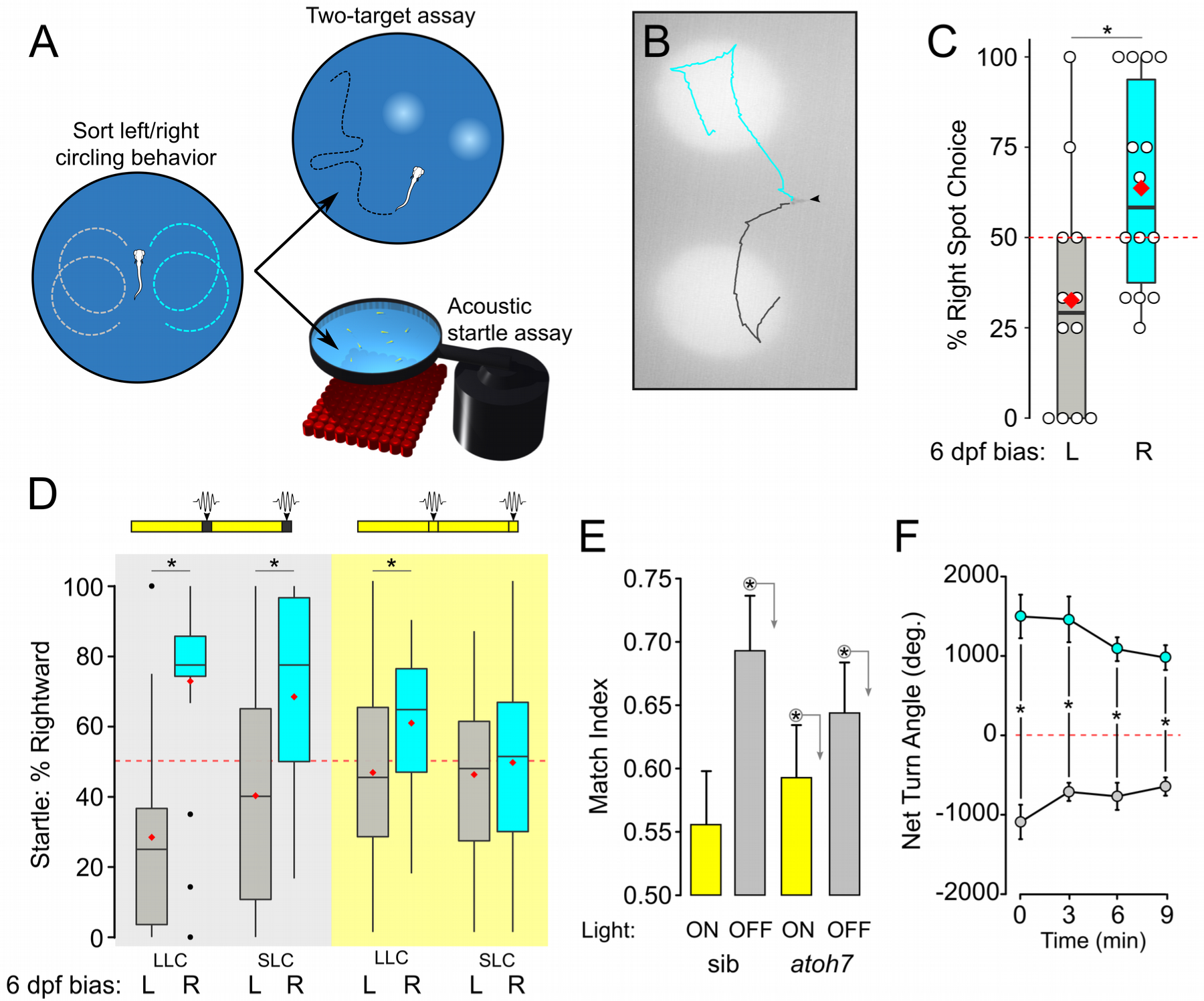
Motor asymmetry is correlated across multiple behavioral tasks. A. Experimental paradigm: larvae were are classified as left or right biased at 6 dpf based on their circling response after loss of illumination (over 4 trials). The following day, the same larvae were tested in either a two-target phototaxis assay or acoustic startle assay. B. Representative path trajectories of individual right (cyan) and left (grey) classified larvae (arrowhead) presented with two symmetric light spots following the loss of illumination. C. Percent of trials on which larvae turned toward the right spot, for larvae classified at 6 dpf as left-biased (grey, N=12) and right-biased (cyan bar, N=13). Each larva performed 4 trials, with individual trials excluded if the larvae was adjacent to the edge of the arena when the phototaxis spots were presented. * p < 0.05, Mann-Whitney U test. D. Percentage of startle responses made in a rightward direction for larvae classified as left (grey) and right (cyan) biased. Larvae were tested either in the dark (grey background) or the light (yellow background). As acoustic stimuli elicit either short or long latency C-starts (SLC, LLC) that are mediated by different circuits, response types were analyzed separately. Red diamond indicates mean. Dark LLC responses: left N=19, right N=20. Dark SLC responses: left N=12, right N=14; Light LLC responses: left N=27, right N=23; Light SLC responses: left N=18, right N=28. * p < 0.05, Mann-Whitney U test. E. Match index for *atoh7* mutants and siblings during baseline illumination (sib, N=45; mutant, N=57; yellow bars) and dark responses (sib, N=51; mutant, N=58; grey bars). * p < 0.05, one-sample permutation test to 0.5. F. Net turn angle for each of four trials after unilateral enucleation of the left (grey, N=4) or right (cyan, N=6) eye. Dotted red line shows random output. * p < 0.05 between groups.

In all three assays, motor asymmetry was primarily manifest in the dark, suggesting that that visual cues over-ride intrinsic bias. We tested this idea using *atoh7* mutants which completely lack projections from the retina to the brain. Even without retinal signaling, zebrafish respond to changes in whole-field illumination via deep brain photoreceptors (Fernandes et al., 2012; Horstick et al., 2017; Kokel et al., 2010). Thus on trials 2-4 of the dark-induced circling assay, both *atoh7* mutants and siblings demonstrated a significant tendency to swim in the same direction as in trial 1 (Figure 2E, grey bars). However, unlike wildtype sibling larvae, *atoh7* mutants also showed a weak but significant motor asymmetry under illuminated conditions (p=0.003; Figure 2E, yellow bars). Next, we used acute unilateral enucleation to control the source of visual information. After acute unilateral enucleation, circling was performed in the direction contralateral to the intact eye on all four trials (Figure 2F). Together, these results support the idea that visual information, when present, predominates over intrinsic motor asymmetry.

### Neurons in the posterior tuberculum maintain left/right motor identity

We next set out to identify the underlying neuronal substrates for left/right bias. Whereas the parapineal and habenula show consistent population-wise anatomical asymmetries, no brain regions are known in zebrafish to have stochastic hemispheric differences in size or function that would be consistent with individual left/right motor bias (Concha et al., 2000; Gamse et al., 2003). We therefore performed a circuit-breaking screen to identify neuronal substrates, crossing Gal4 lines that label distinct brain structures to a UAS:epNTR-TagRFP reporter for cell-specific ablation using nitroreductase (Figure 3A) (Horstick et al., 2014; Pisharath et al., 2007). After lesioning the labeled population of neurons in each line, we then tested whether individual motor bias remained. Larvae showed a decrease in motor bias after ablation of neurons in two Gal4 lines: *y279* and *y375* (Figure 3B). Reduced motor bias was specific, as ablation did not affect the total amount of turning behavior or baseline locomotion (Figure S2A). Finally, the chemogenetic ablation protocol itself did not disrupt motor bias as ablation of neurons in orthopedia driver line *otpb.A:Gal4* had no effect (Figure 3B). Thus the *y279* and *y375* Gal4 lines label neurons that are critical for individual motor bias.

**Figure 3:**
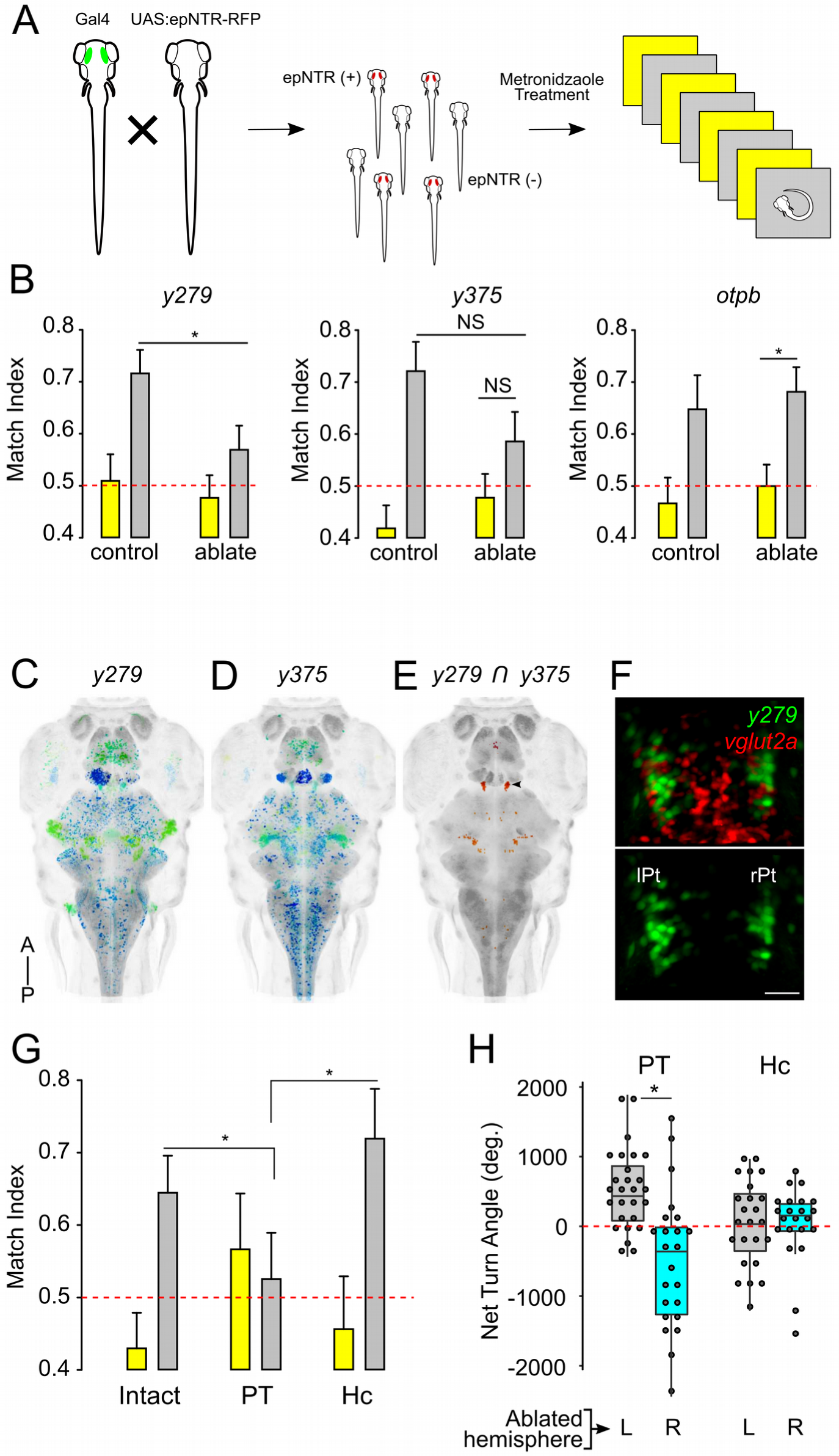
Neurons in the posterior tuberculum maintain left/right identity. A. Chemogenetic ablation screen: transgenic lines with restricted Gal4 patterns were crossed to a UAS:epNTR-RFP reporter. Both epNTR-RFP+ and non-fluorescent siblings (as controls) were treated with metronidazole before testing for motor asymmetry under light and dark conditions. B. Match index for drug treated controls (*y279* N=54; *y375* N=43; *otpbA* N=35) and following genetic ablation (*y279* N=58; *y375* N=37; *otpbA* N=46) during paired baseline (yellow) and dark (grey) responses. C-D. Whole brain dorsal Zebrafish Brain Browser (ZBB) projections for *y279* (D), *y375* (E). Color is depth scale. E. Computed intersect *y279* and *y375* expression patterns. Arrowhead indicates PT. F. Dorsal confocal projection through the rostral PT in *y279-Gal4, UAS:Kaede* (green) crossed to *vglut2a:dsRed* (red). Scale bar 20µm. G. Match index in unablated controls (N=45) and after bilateral laser ablation of the PT (N=17) and Hc (N=19) during baseline (yellow) and on dark trials (grey). * p < 0.05, Mann-Whitney U test. H. Net turn angle (mean on trials 1-4) for left PT hemisphere (grey, N=27) and right PT hemisphere (cyan, N=23) ablations. Hc unilateral ablations (right bars) (left hemisphere grey, N=24; right hemisphere cyan, N=22). * p < 0.05, t-test.

*y279* and *y375* each express Gal4 in neurons distributed in multiple brain regions (Figure 3C,D). We reasoned that clusters labeled by both lines would be strong candidates for driving lateralized behavior. Because both lines express the same reporter, we could not directly identify co-labeled neurons, but instead compared co-registered whole-brain images of each line (Marquart et al., 2015). Salient virtual co-expression was present in a bilateral cluster of neurons in the posterior tuberculum (PT) (Figure 3E, S2B-E). Cell counts established that each PT hemisphere comprised 28.3 ± 7.2 *y279^+^* cells (mean ± standard deviation, N=28 larvae, Figure 3F), but did not reveal differences between hemispheres in left and right biased larvae (Figure S2F). We assessed whether the PT clusters were required for left/right bias by using focal laser ablation of *y279* neurons. As a control we also ablated the caudal hypothalamus (Hc) which is strongly labeled in *y279*. Ablation of the Hc had no effect whereas bilateral ablation of the rostral PT cluster strongly reduced motor bias, consistent with a loss of left/right identity (Figure 3G, S3A). Conversely, after unilateral PT ablations 76% of larvae (38/50) circled in the direction ipsilateral to the intact PT during dark trials, with no effect during baseline illumination (Figure 3H, S3B-C). The posterior tuberculum is a ventral region of the diencephalon that includes identified classes of dopaminergic (DA) neurons (Rink and Guo, 2004; Tay et al., 2011). However, the cluster of PT neurons labeled in *y279* and *y375* is situated rostral and dorsal to PT DA neurons (Figure S2B). Moreover *otpa* mutants, which lack key classes of DA neurons, and *otpb.A:Gal4* ablations retained robust left/right bias (Figures 3B; S3D). It is therefore unlikely that PT dopaminergic function contributes to motor asymmetry. These results confirm that a cluster of neurons in a rostral lobe of the posterior tuberculum impose left/right bias on motor responses in zebrafish.

### PT neurons show persistent activity after loss of illumination

We next examined whether PT neurons are active after loss of illumination by imaging changes in GCaMP6s fluorescence in *y279-Gal4, UAS:GCaMP6s* larvae. Because we previously found that photic stimulation acutely terminates localized circling behavior (Horstick et al., 2017), we used an infra-red laser for two-photon excitation of GCaMP6s during these recordings. We attempted to simultaneously record tail movements, but found that persistent motor responses to the loss of illumination were not apparent in immobilized larvae. Nevertheless we reasoned that even if the motor behavior was not preserved, PT neurons might still respond to loss of illumination. Indeed, 16.2% of PT neurons (37/228 from 6 larvae) responded to the light off stimulus (Figure 4A-B). PT neuron activity peaked shortly after loss of illumination and remained elevated during the 57s recording interval, consistent with the time-frame of circling behavior (Figure 4C). An additional 3.9% of PT neurons (9/228) were active upon the restoration of illumination. However no significant correlations were observed between the left/right identity of individual larvae and peak GCaMP activation or position of light OFF responsive neurons within the PT (Figure 4D,E). These results establish that a subset of PT neurons fire in response to changes in illumination and remain active on a time scale consistent with the duration of motor asymmetry during dark-induced circling behavior.

**Figure 4:**
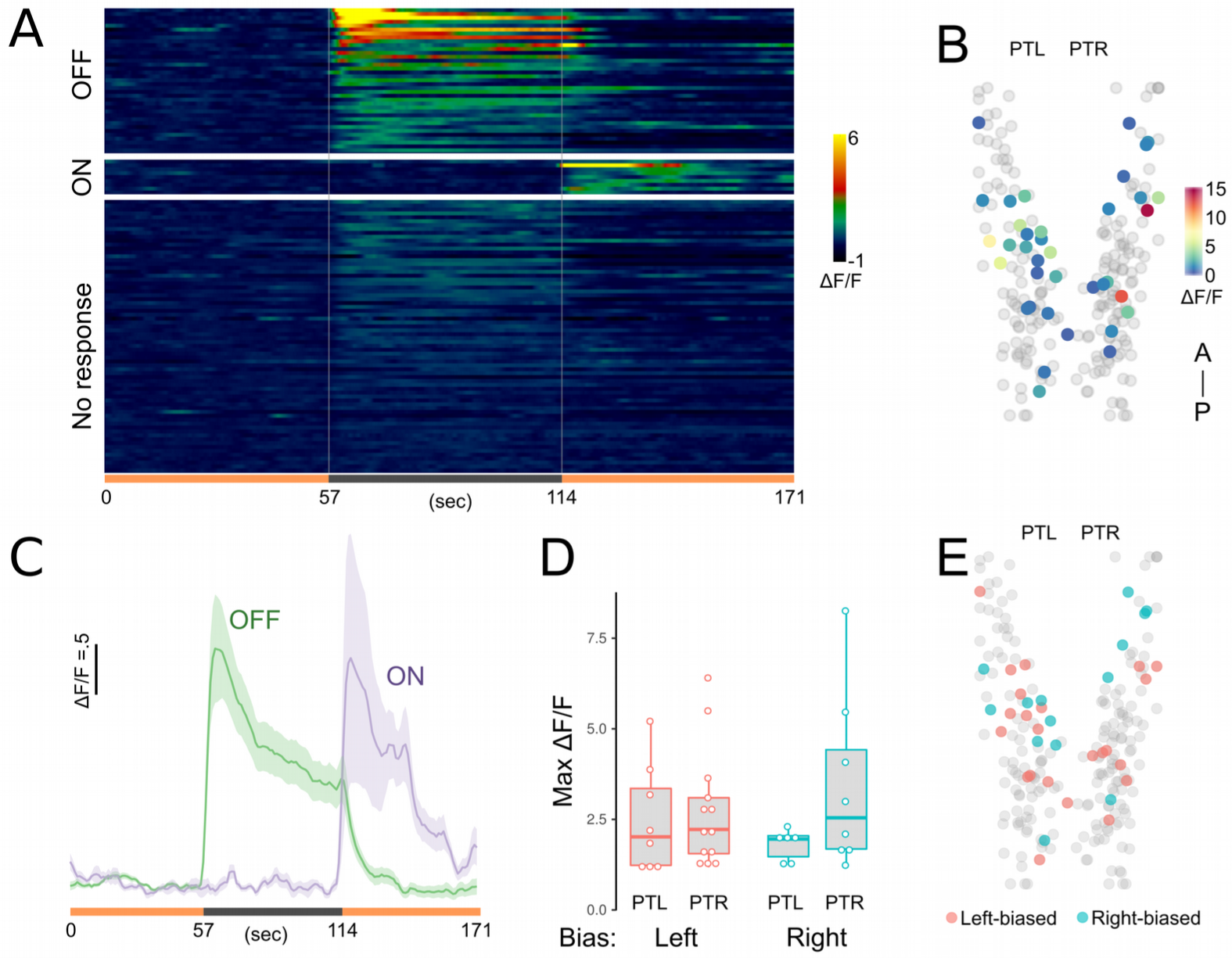
Neurons in the posterior tuberculum respond to changes in illumination. A. Raster plot of mean calcium responses (mean of 3 trials) from GCaMP6s expressing PT neurons. Color scale denotes standardized change in fluorescence intensity (ΔF/F). Illumination conditions as indicated on x-axis (Light ON, orange; dark, grey). Only a subset of no response neurons are included. B. Location of light OFF responsive neurons in the rostral PT. Scale bar indicates fluorescence (ΔF/F) change over baseline. C. Mean and standard error for response of light OFF (green, N=37) and light ON (purple, N=9) responsive neurons. D. Peak change in fluorescence for neurons that respond to light OFF in the left (PTL) and right (PTR) PT for larvae classified as left (red) and right (blue) motor biased. E. Location of light OFF responsive neurons within the PT for larvae behaviorally identified as left (red) or right (blue) biased.

### Projections from the PT to habenula are essential for motor bias

The epithalamic commissure (Ec) that runs between the habenula hemispheres is labeled in *y279-Gal4*, *UAS:Kaede* larvae (Figure 5A). As *y279* also labels neurons in the habenula — particularly in the left habenula — we initially suspected that this commissure was formed by *y279* habenula neurons. However when performing confocal scans to verify laser ablations, we noticed that fluorescently labeled fibers were absent in bilateral PT lesioned larvae, raising the possibility that commissural fibers originate in the PT (Figure 5B). To test this, we unilaterally photoconverted Kaede in the PT in *y279-Gal4, UAS:Kaede* larvae. Photoconverted Kaede labeled neurites that emerged dorsally from each PT hemisphere and projected to the ipsilateral habenula. Within the habenula these fibers turned and crossed to the contralateral side through the epithalamic commissure (Figure 5C,D). PT projections terminated in the neuropil region of both habenula hemispheres. No salient differences in the PT-habenula projection were observed for left or right-biased larvae, however this observation raised the possibility that the PT controls motor bias via signaling to the habenula. We tested this by laser ablation of *y279* habenula neurons (Figure S3A). Bilateral ablation of *y279* habenula neurons abolished left/right bias without affecting total turning (Figure 5E-F). Moreover after unilateral habenula ablation motor responses were strongly lateralized during dark-induced circling behavior, without inducing motor bias under baseline illumination (Figures 5G; S3C). In contrast to PT lesions, total turning behavior did not increase after unilateral habenula ablation (Figure S3E). Intriguingly, selective lesion of the epithalamic commissure also induced a strong motor asymmetry that was similar to ablation of the right-habenula, such that larvae became left-biased only under dark conditions (Figure 5G,H, S3C,F). Together, the loss of motor bias after bilateral ablation of either the PT or habenula, induction of left/right lateralized behavior after unilateral ablations and presence of a direct connection between these areas, argues that the PT-habenula pathway imposes left/right bias on motor responses.

**Figure 5:**
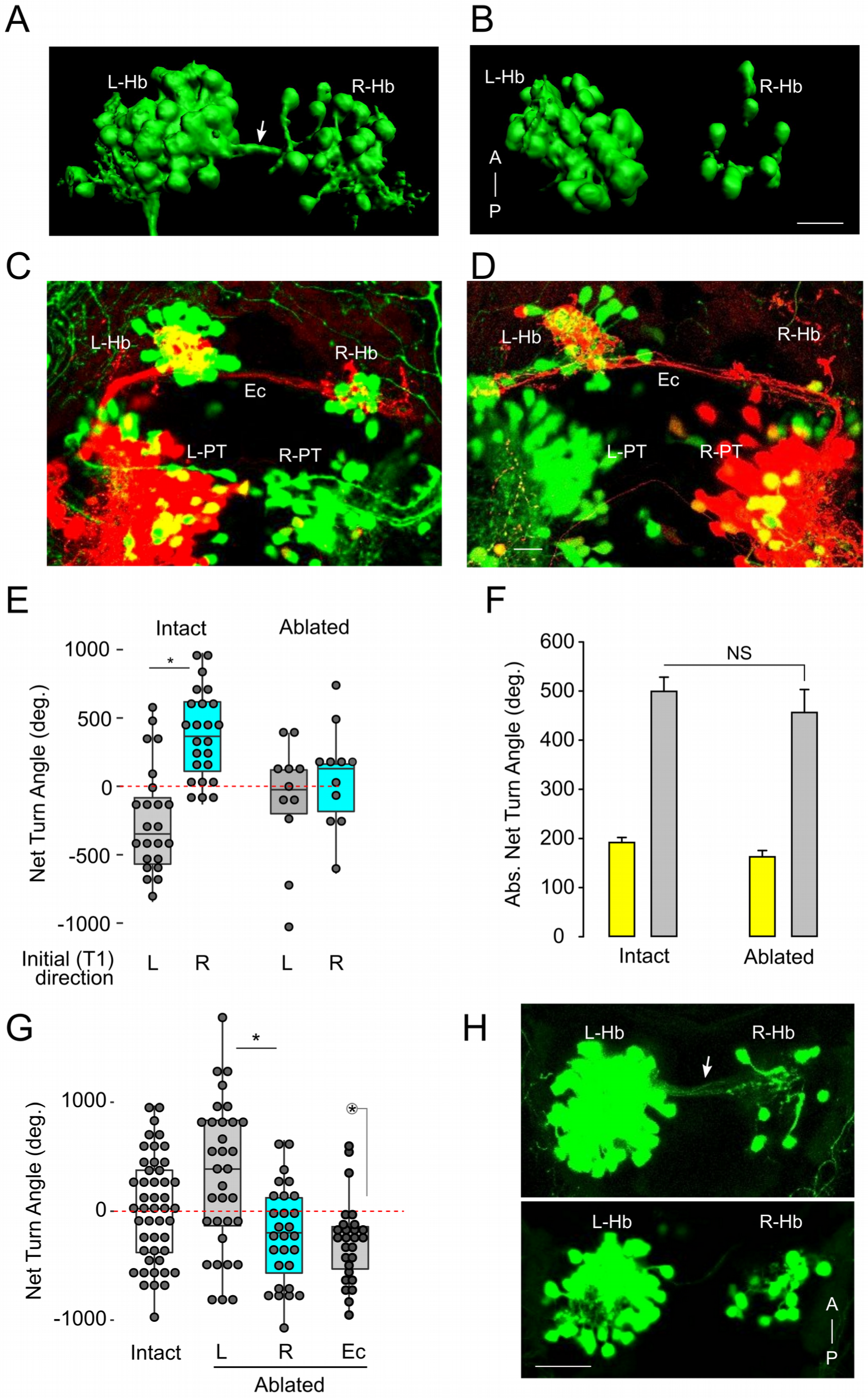
A projection from posterior tuberculum to habenula drives motor asymmetry. A-B. 3D rendering of *y279-Gal4, UAS:Kaede* expression in the habenula at 7 dpf in intact larvae (A) and after bilateral ablation of *y279* expressing PT neurons (B). Arrow indicates epithalamic commissure (Ec) present in intact larvae that is lost after bilateral PT ablation. Scale bar 20µm. C-D. Confocal projections of *y279-Gal4;UAS:Kaede* expressing larvae following unilateral focal photoconversion of Kaede in the left (C) or right (D) rostral PT. Photoconverted Kaede in the red channel is saturated to facilitate visualization of PT projections. Scale bar 20µm. E. Net turn angle (mean of trials 2-4) for intact control larvae (N=47) and after bilateral ablation of *y279* expressing habenula neurons (N=22). Larvae were classified as left-biased (grey) or right-biased (cyan) based on trial 1 (T1). * p < 0.05, t-test. F. Total amount of turning for larvae in (E) during locomotion under baseline illumination (yellow) or dark-induced circling behavior (grey). G. Net turn angle (mean of trials 1-4) in non-ablated controls (white, N=47) and following unilateral ablation of the left (grey, N=33), right (cyan, N=28) habenula nuclei or following laser section of the epithalamic commissure (Ec) (N=24). * p < 0.05 t-test. circled * p < 0.05, one-sample t-test to 0. H. Representative confocal scan showing *y279* labeled epithalamic commissure (Ec) (arrow) in a control larva (top). Following ablation commissure is absent (bottom). Scale bar 20µm.

### Haploinsufficient notch pathway mutations disrupt lateralized behavior

Many species, including zebrafish, show a stereotyped asymmetric development of specific brain nuclei and placement of viscera. Genes involved in this process have been relatively well characterized (Concha et al., 2000; Lopes et al., 2010; Yan et al., 1999). However much less is known about genetic pathways that lead to the stochastic acquisition of left or right motor asymmetry. During the course of our studies, we isolated a background mutation (*y606*) in our wildtype stock that weakened motor bias during circling behavior. In affected clutches, 25% of embryos showed a ‘curly-up’ phenotype that severely effected tail morphology and precluded behavioral testing (Figure 6A). We speculated that the morphological abnormality represented a homozygous phenotype, with loss of motor asymmetry present in morphologically normal heterozygous larvae (Figure 6B). RNAseq-based bulk segregant mapping linked the mutation to a six Mbp interval on chromosome 9 and revealed a 7.7-fold reduction in the expression of the gene *epb41l5* in this interval in *y606* mutants compared to siblings (Figure S4A-B) (Miller et al., 2013). No RNAseq reads mapped to the first two exons, and we were not able to amplify these exons from mutants using genomic PCR (Figure S4C). Consistent with this, genomic PCR revealed a 4.4 kb deletion that excises exons 1 and 2 of *epb41l5*, eliminating the transcription and translation start sites (Figure S4D). *Epb41l5* is the mutated gene in the *mosaic eyes* (*moe^b476^*) mutant that shows a similar curly-up phenotype (Jensen et al., 2001) and *moe^b476^* failed to complement *y606*, confirming that y606 is a new allele of mosaic eyes.

**Figure 6:**
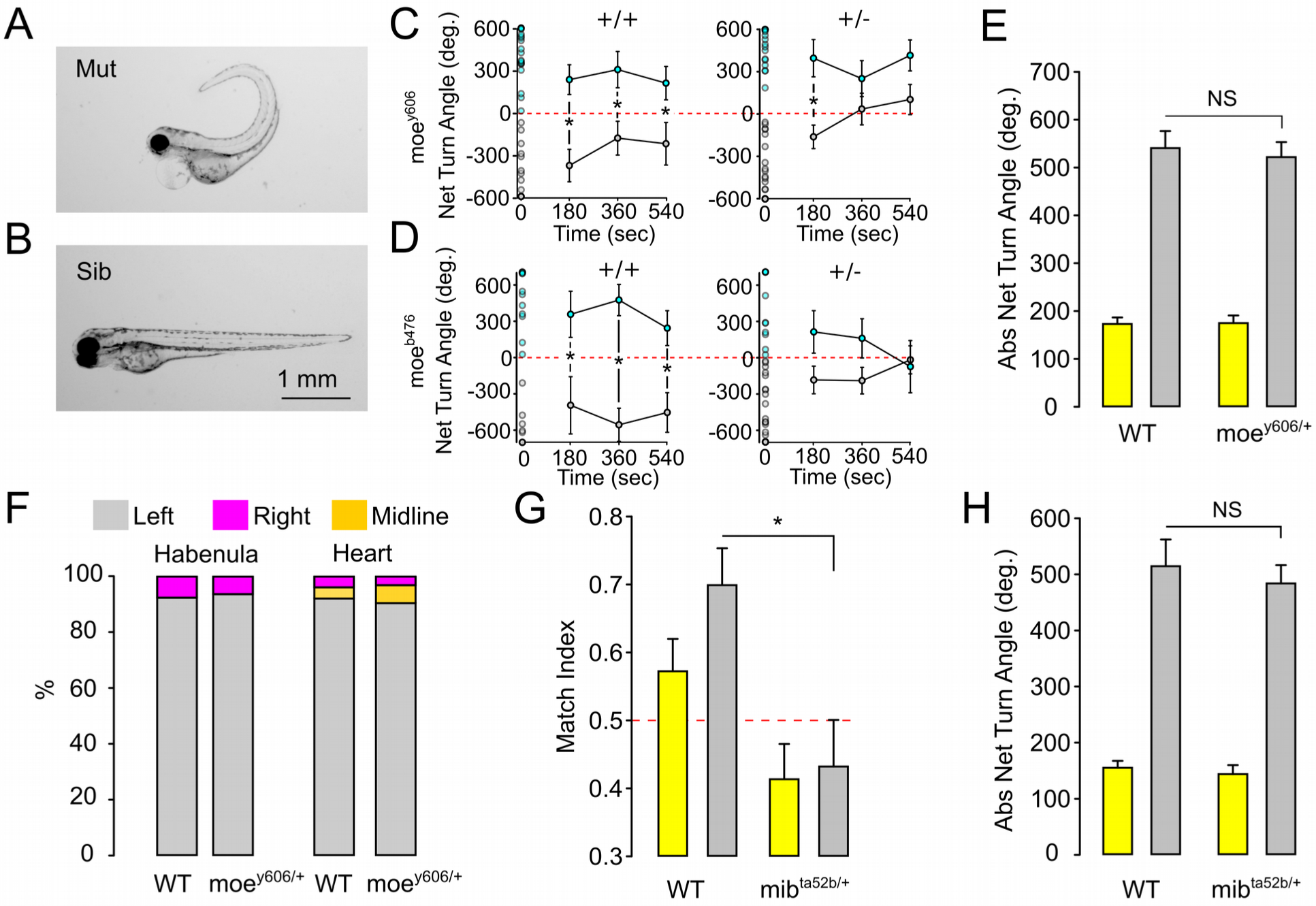
Left/right identity is disrupted by mutations in genes that regulate Notch signaling. A-B. Curly-up morphology in 2 dpf *y606* mutants (A) and sibling larvae (B). C. Net Turn Angle for *moe^y606^* heterozygous (left, +/-) and wildtype (right, +/+) sibling larvae over four 30 s light off trials. Open circles in the first trial (time 0) represent individual NTA for all larvae tested and were used to classify larvae as right (cyan, Het N=34; WT N=20) or left-biased (black, Het N=27; WT N=20). Subsequent points represent mean for left/right groups on trials 2-4. D. Same analysis as in (C) for *moe^b476^* allele showing right (cyan, Het N=11; WT N=15) or left-biased (black, Het N=25; WT N=9) larvae. * p < 0.05 between groups in C-D. E. Absolute Net Turn Angle for moe^y606^ heterozygous (N=47) and wildtype sibling (N=40) larvae averaged over 4 trials for baseline illumination (yellow bar) and light off (grey bar) trials. F. Habenula and heart placement in wildtype sibling and moe*^y606^* heterozygous larvae. Left: Percentage of larvae with larger habenula hemisphere on each side (N=28, 33 for wildtype, hets). Right: Percentage of embryos with heart positioned on the left, right or midline (N=28,34 for wildtype, hets). G-H. Match index (F) and total turning (G) for wildtype sibling and *mib^ta52b^* heterozygous larvae (N=31,27 respectively), during baseline (yellow) and dark (grey) responses. * p < 0.05, Mann-Whitney U test.

We incrossed *moe^y606^* heterozygous adults and tested morphologically normal siblings of curly-up mutants in the dark-induced circling assay, then performed post-hoc genotyping to distinguish wildtype and heterozygous larvae. Wildtype larvae showed normal motor asymmetry, however heterozygous *moe^y606^* larvae showed a significant reduction in left/right-bias (Figure 6C). Other motor parameters and asymmetric development of the epithalamus and viscera were unaffected in heterozygotes (Figure 6E,F). To corroborate these findings we used *moe^b476^*, an independent allele that we confirmed eliminates the entire *epb41l5* gene (Figure S5A-C). Heterozygous mutations in *moe^b476^* also disrupted motor bias during circling behavior (Figure 6D). These results demonstrate that acquisition of left/right identity in zebrafish requires two functional alleles of *epb41l5* in zebrafish.

*Epb41l5* regulates the timing of neurogenesis, interacting with proteins in the *Notch* signaling pathway (Matsuda et al., 2016; Ohata et al., 2011). Notch regulates *Nodal*, the central gene that drives the development of molecular and anatomical left-right asymmetry (Krebs et al., 2003; Raya et al., 2003). We therefore reasoned that other heterozygous mutations in Notch signaling proteins might perturb left/right bias and tested larvae with heterozygous mutations in *mindbomb* (*mib*), an E3 ubiquitin ligase that is essential for notch signaling (Itoh et al., 2003). Indeed, *mib* heterozygotes showed a severe loss of left/right-bias during dark-induced circling (Figure 6G) without perturbing the total amount of turning (Figure 6H). Together, these findings reveal that acquisition of left/right motor identity is disrupted by mutations in genes that regulate Notch signaling levels during embryonic development.

## Discussion

Here, we reveal that in the absence of visual information, zebrafish manifest a durable left/right motor bias that is driven by neurons in the posterior tuberculum. Lateralized motor behavior occurs both during spontaneous movement, and in response to visual and auditory cues. Zebrafish do not show a population-level bias, unlike in humans where 90% of the population favor the right hand (Corballis, 2003). Rather, similar to motor asymmetries in many other species, lateralized behavior is manifest as an individual bias to execute movements in a preferred direction on ~70% of trials. Left/right identity is maintained over at least several days despite handling and changes in the environment. Bilateral clusters of ~30 genetically identified neurons per hemisphere in the rostral lobe of the posterior tuberculum are essential for the expression of this motor asymmetry. This conclusion is supported by loss of lateralized behavior after chemogenetic ablation of rostral PT neurons using two transgenic Gal4 lines that have minimal overlap elsewhere in the brain, and direct laser photoablation of the PT. In addition, unilateral laser ablation of PT neurons was sufficient to create a population of fish whose response direction was heavily biased to the side of the intact PT. Critically, after rostral PT lesions, larvae continued to show persistent turning to one side after loss of illumination, but circling behavior was initiated in a random direction on each trial rather than in a preferred direction. It is therefore unlikely that the rostral PT is part of the sensory pathway that initiates dark-induced circling, but rather suggests that the PT is a locus that maintains left/right identity and imposes this bias on motor responses. Finally, neurons in the rostral PT show sustained activity after loss of illumination, consistent with the duration of lateralized circling movements. Together, these findings confirm that the rostral posterior tuberculum drives lateralized behavior in zebrafish and define a specific neuronal substrate for a lateralized motor behavior in a vertebrate.

### A PT-habenula circuit directs motor asymmetry

Our findings suggest a model in which bilateral PT-habenula units compete to strengthen, but not directly drive ipsilateral dark-induced circling behavior (Figure 7). In individual larvae, one hemisphere predominates, providing a greater drive to premotor circuits. How left/right identity is encoded in the PT-habenula unit is unclear: we did not detect differences in the number of neurons or light-off activity between hemispheres that correlated with motor bias. The strong effect of unilateral PT ablations indicate that the intact hemisphere, even if it was not previously dominant, was capable of imposing ipsilateral motor bias, suggesting that dominance is unlikely to reflect a unique quality present in only one hemisphere. However, after unilateral PT ablation, not only was dark-induced motor behavior strongly lateralized, but net turning also increased, consistent with the idea that in the absence of competition, there is greater drive biasing ipsilateral premotor circuits. Although we are not aware of evidence for a direct connection between PT clusters we found that PT neurons project to and commissurally within the habenula, concordant with a previous report (Hendricks and Jesuthasan, 2007). Thus competition between PT clusters may take place within the habenula. In keeping with this idea, ablation of *y279* habenula neurons disrupted lateralized responses similar to PT ablation. However, ablation of the habenula commissure yielded a population of larvae with left-biased responses, similar to ablation of either right habenula or PT, possibly suggesting that the left habenula predominates in the absence of communication. Thus, although our findings do not resolve the precise mechanism by which the PT directs motor asymmetry, they demonstrate that a PT-habenula pathway is an essential substrate for lateralized behavior in zebrafish.

**Figure 7:**
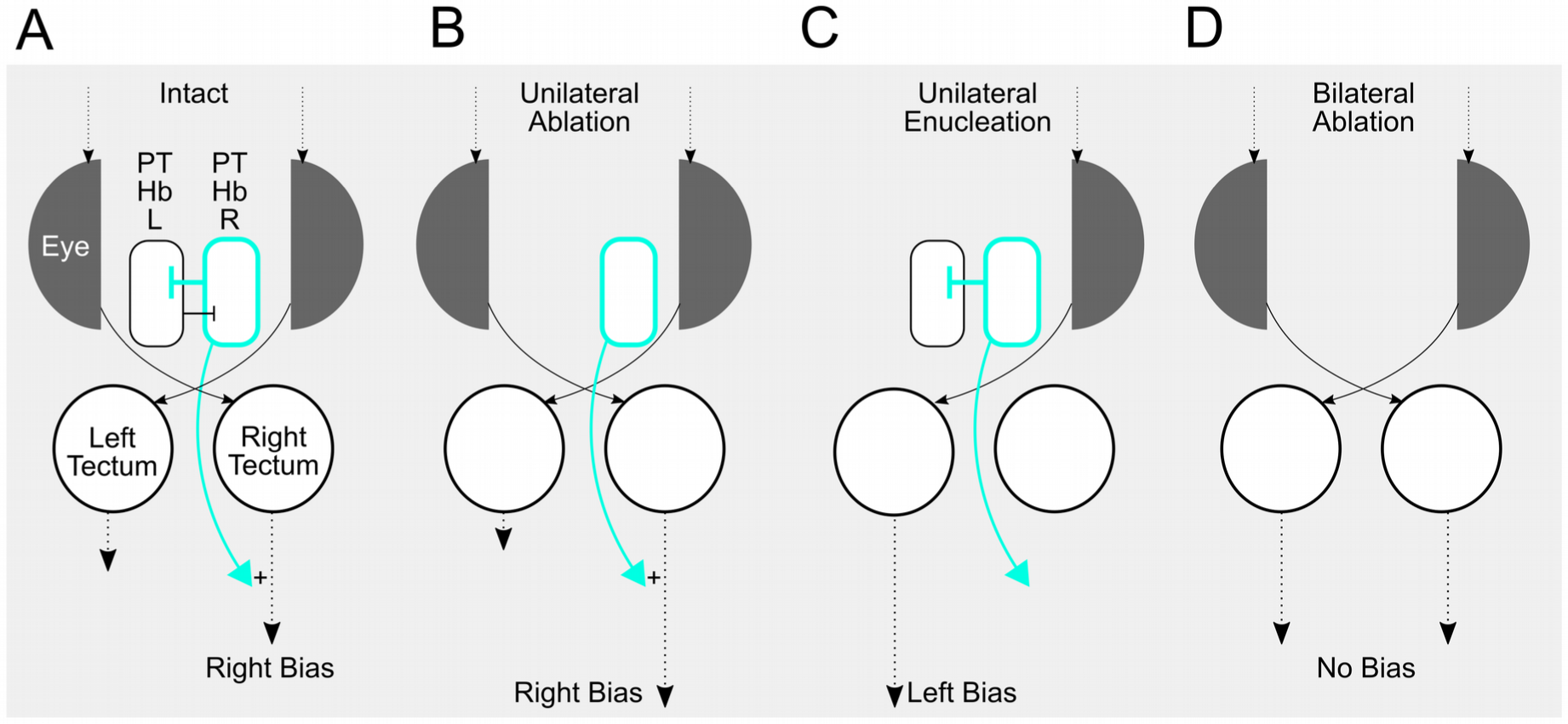
Motor asymmetry in zebrafish. A. For a right biased larva, the dominant right PT-habenula pathway imposes right motor bias by modulating symmetric visual drive and suppressing the left PT-habenula pathway. B. Competition between PT-habenula units is eliminated after unilateral ablations, strengthening motor bias. C. Asymmetric visual input after unilateral enucleation overrides PT-habenula modulation of motor output. D. Symmetric visual stimuli, after bilateral ablation of PT-habenula units eliminates motor bias leading to randomized turn direction.

### Genetic control of individual left/right identity

The last two decades has produced significant progress in understanding the morphogenetic processes that lead to consistent left-right asymmetries within the nervous system (Roberson and Halpern, 2018). In contrast, much less is known about how stochastic individual patterns of left/right identity are established. In birds, asymmetric sensory experience during a critical period in early development imprints hemispheric dominance for visual feature processing (Koshiba et al., 2002; Rogers, 1982). In contrast, we found that dark-reared larvae and *atoh7* mutants which lack central projections of retinal ganglion cells at all stages maintained individual motor lateralization indicating that neither spontaneous retinal activity nor asymmetric visual experience are required for individual bias to emerge. Further, as in mice and *Drosophila*, selective breeding experiments excluded the possibility that a heritable factor dictates left/right preference (Figure S6)(Buchanan et al., 2015; Collins, 1969). Most likely, directional bias is stochastically determined during embryogenesis by either a specific symmetry breaking event or by natural variability in development. While the nature of this process is not yet known, recent genome-wide association studies have provided tantalizing hints that molecular pathways involved in the development of visceral asymmetries also contribute to the establishment of human handedness (Brandler et al., 2013). Consistent with this idea, we found that left/right motor identity was disrupted by heterozygous mutations in *epb41l5*, a gene required for left-right patterning in mice (Lee et al., 2010). *Epb41l5* is a membrane adapter protein (Matsuda et al., 2016; Ohata et al., 2011) that interacts with *mind-bomb*, a ubiquitin-ligase required for Notch signaling during neurogenesis (Itoh et al., 2003). Accordingly we found that heterozygous mutations in *mind-bomb*, a ubiquitin-ligase required for notch signaling, also disrupted lateralized behavior. Surprisingly, behavioral changes were seen in haploinsufficient mutants suggesting that acquisition of left/right identity is highly sensitive to notch signaling levels. Indeed in mice, haploinsufficient mutations of the notch ligand *delta-like 1* lead to a reduction in dopaminergic neuron differentiation, and in zebrafish manipulations that augment or disrupt notch signaling during habenula neurogenesis can shift the production of lateral or medial neurons and thereby isomerize the habenulae (Aizawa et al., 2007; Trujillo-Paredes et al., 2016). Intriguingly, *notch1a* has a similar temporal profile of expression in the developing PT raising the possibility that heterochronic neurogenesis of PT neuron subtypes is involved in acquisition of left/right identity (Mueller and Wullimann, 2005). Thus although we did not observe changes in the number of *y279* expressing neurons in the left or right PT hemisphere in *epb41l5* heterozygous mutants (Figure S7), future studies may reveal alterations in the differentiation of neuronal cell types associated with loss of left/right identity.

### Neural correlates of motor asymmetries

Cerebral asymmetries have been proposed to increase information processing power through hemispheric specialization (Vallortigara and Rogers, 2005), or, to reflect space constraints in the brain that necessitate partitioning processing functions between hemispheres (Barneoud and Van der Loos, 1993). Indeed, the left and right habenula nuclei in zebrafish are activated by different sensory modalities and have separable roles in behavior (Cheng et al., 2017; Dreosti et al., 2014; Duboue et al., 2017; Krishnan et al., 2014; Zhang et al., 2017), and in *C elegans* and *Drosophila* brain asymmetries have been linked to sensory processing and memory formation respectively (Pascual et al., 2004; Pierce-Shimomura et al., 2001; Wes and Bargmann, 2001). Asymmetries in sensory processing areas in birds are correlated with lateralized control of behavior (Rogers, 1990). Similarly, it seems natural to postulate that asymmetric structural or functional properties of the nervous system also underlie motor asymmetries. However, evidence for this idea is surprisingly elusive: motor lateralization has only rarely been correlated with an underlying cerebral asymmetry, and causal relationships have yet to be established (Davison et al., 2009; Gutierrez-Ibanez et al., 2011; Jozet-Alves et al., 2012; Lee et al., 2017). The strongest relationship to date has been described in the pond snail *Lymnaea stagnalis* where the direction of coiling behavior by males during mating almost completely matches the side on which two central ganglia are fused (Davison et al., 2009).

Even less clear is the nature of the ethological advantage conferred by motor asymmetries. An enduring argument is that the hemispheres of an ideal perfectly symmetrical nervous system can not discriminate bilaterally symmetrical sensory stimuli and would therefore reach a stalemate in selecting a left or right motor response to a non-directional cue (Webster, 1977). Intrinsic neural asymmetries may therefore facilitate motor responses to stimuli with ambivalent directional information, avoiding simultaneous initiation of left/right responses, or reactions that are delayed due to difficulty selecting a response direction. Indeed, during dark-induced circular swimming, larvae must initiate movement by contracting muscles on one side. In the light, even small asymmetries in the visual environment may over-ride innate bias, explaining why we do not observe motor asymmetry under illuminated conditions. This idea is also consistent with our finding that *atoh7* mutants, which are blind, show motor bias under both light and dark conditions; in mutants visual stimuli do not reach the brain to over-ride innate direction preference.

Motor asymmetries such as human handedness are among the most pervasive and salient forms of individual variation. Moreover, variation in motor asymmetries is linked to inter-individual differences in personality, cognitive processing and risk for neurodevelopmental disorders (Markou et al., 2017; Sommer et al., 2001). Yet, attempts to understand how such motor asymmetries are stochastically generated during development, or even their structural or functional basis in the brain, have yielded limited insight. In particular, efforts to study the development and basis for human handedness have been hampered by difficulty in identifying a neural substrate, and inability to perform experimental manipulations. Our identification of a specific neuronal substrate for asymmetric motor behavior in zebrafish opens up a new model for understanding how functional lateralization emerges and is maintained in the nervous system. Moreover our data also suggests that acquisition of left/right identity is sensitive to specific levels of *Notch* pathway activity. These results therefore set the stage to uncover precisely how motor bias is established during development and is maintained by structural or functional asymmetries within the nervous system.

## Acknowledgements

We thank Jennifer Sinclair for expert technical support, Dr. Katie Drerup for assistance with RNAmapper, Greg Palardy for the mib genotyping protocol and the NICHD Molecular Genomics Core for library preparation and sequencing. This work was supported by the Intramural Research Program of the *Eunice Kennedy Shriver* National Institute for Child Health and Human Development (NICHD) and utilized the high-performance computational capabilities of the Biowulf Linux cluster at the National Institutes of Health, Bethesda, MD.

## Author Contributions

EJH and HAB conceived the experiments and wrote the manuscript. EJH, YB and HAB performed experiments. All authors approved the final manuscript.

## Methods

### Zebrafish

Adult zebrafish (*Danio rerio*) were maintained with a Tubingen long fin strain background. All animal care and experimental procedures were approved by the NICHD animal care and use committee. Experiments were performed on larvae in the first 10 days post fertilization (dpf), before sex differentiation. Larval zebrafish were raised on 14/10 h light/dark cycle at 28 °C, at a maximum density of 20 in 10 mL E3h medium (5 mM NaCl, 0.17 mM KCl, 0.33 mM CaCl2, 0.33 mM MgSO4, 1.5 mM HEPES, pH 7.3). For dark-rearing at 3 hours post-fertilization embryos were sorted into 60 mm Petri dishes at a stocking density of 15 larva per 10 mL medium and placed into a dark box till 5 dpf. At 5 dpf the media was replaced and larvae maintained under normal light cycles till testing at 6 dpf. For controls, siblings were raised in parallel under the same conditions except with exposure to normal 14/10 light cycles.

Transgenic lines used were enhancer traps *y279-Gal4* and *y375-Gal4* (Marquart et al., 2015), *Tg(UAS:epNTR-tagRFP)y268* (Tabor et al., 2014), *Tg(UAS:Kaede)s1999t* (Davison et al., 2007), *Tg(atoh7:GFP)rw021* (Masai et al., 2003), *Tg(otpb.A:Gal4-myl7:GFP)zc67* (Fujimoto et al., 2011) and *TgBAC(vglut2a[slc17a6b]:loxP-mCherry-loxP-Gal4ff)nns21* (Satou et al., 2013). Mutant lines: *otpa^m866^* (Fernandes et al., 2012). *moe^b476^* and *mib^ta52b^* were kind gifts of Ajay Chitnis (Jensen et al., 2001; Schier et al., 1996). Gal4 lines are available from the Zebrafish International Resource Center (http://zebrafish.org).

### Behavior tracking and analysis

Behavioral tests were performed on 6 and 7 dpf larvae except as noted. Infrared illumination (CMVision Supplies, 850 nm) was used to monitor larvae with camera lenses fitted with IR-longpass filters to exclude visible light. Visible illumination (40 µW/cm^2^ measured using a radiometer (International Light Technologies)), was provided by white LEDs positioned over the recording chamber (Thorlabs). Testing areas were maintained at 26-28°C and larvae were adapted to the recording environment for 30 min prior to starting experiments. We used DAQtimer event control software to coordinate illumination conditions and recordings (Yokogawa et al., 2012). When experiments required confocal brain imaging and behavioral analysis, larvae were raised in medium containing 200 µM PTU starting at 1 dpf. PTU was removed at least 24 h prior to behavioral recordings.

#### Trajectory Analysis

Path trajectory recordings were performed and analyzed as previously described (Horstick et al., 2017): images were captured with a uEye IDS1545LE-M CMOS camera (1^st^ Vision) and larvae tracked in real-time using DAQtimer. Individual larvae were placed into a 120 x 120 mm arena. Each individual was tracked for 30 s after loss of illumination over 4 successive trials separated by 3 min baseline illumination (with the same protocol used for constant illumination trials except the light remained on throughout). Where behavioral experiments were coupled with manipulations (chemogenetic or laser ablation, genetic mutations) baseline controls were obtained by recording the 30s illuminated interval prior to the light off interval except as otherwise noted. We used three measures to characterize directional bias in path trajectories: net turn angle (NTA), Absolute NTA and Match Index. **NTA**: the sum total of all leftward (- degrees) and rightward (+ degrees) path direction changes over a 30 s recording interval. Thus, individuals that changed direction equally to the left and right would have an NTA of 0, indicating no net directional preference. **Absolute NTA**: as for the NTA but taking the absolute value of each path direction change before summation. The absolute NTA measures the total amount of turning behavior during a 30 s interval. **Match Index**: Used for experiments where larvae were subjected to four trials, the Match Index is the fraction of trials 2-4 where the NTA had the same sign as on the first trial. We excluded trials where we obtained less than 10 s of data for an individual. A MI of 1.0 indicates that a larvae turned in the same direction on trials 2-4, as on trial 1, whereas a MI of 0.33 would occur if only one of trials 2-4 were performed in the same direction as on trial 1.

#### Kinematic analysis

To measure routine turn initiation larvae were recorded using a high-speed camera (DRS Lightning RDT/1; DEL Imaging) at 1000 Hz with off-line analysis of video images using Flote (Harold A Burgess and Granato, 2007). Larvae were tested in a 58 x 58 mm arena with IR illumination and recordings were triggered when larvae entered a centrally placed 15 x 15 mm ROI using DAQtimer software and a uEye camera. To minimize environmental visual cues that might influence trajectories, the arena was completely enclosed with a diffuser underneath the arena and a fitted lid above the arena holding a IR long-pass filter. As for trajectory analysis, we performed 4 recordings (10 s duration) after loss of illumination separated by 3 min of baseline illumination. Control experiments were the same, but without the dark periods. We analyzed only larvae that executed at least 3 routine turns on each of the 4 trials.

#### Startle direction assay

Acoustic startle tests were performed as previously described (Harold A. Burgess and Granato, 2007). To determine if startle direction bias correlated with dark-circling direction bias, larvae were classified as left/right at 6 dpf using trajectory analysis over 4 trials. At 7 dpf larvae were placed into a 3 x 3 grid of 1 x 1 cm wells. For trials in the dark, larvae were tested with 16 repeats of an auditory/vibrational stimulus (~18dB relative to 1 m/s^2^), occurring 10 s after loss of illumination. Each repeat was separated by 2 min constant illumination. Trials in the light were the same, without the 10 s dark period. Only larvae performing at least 4 long or short latency startle responses were analyzed. For analysis we compared the percentage of short and long latency C-starts performed in a rightward direction for each pre-classified group of larvae.

#### Two-target phototaxis assay

For the two-target phototaxis assay, we first identified the left/right identity of larvae at 6 dpf using trajectory analysis over 4 dark trials. Larvae with consistent directional responses to all 4 trials were then retained for testing the next day. At 7 dpf, these larvae were individually placed into a 58 x 58 mm arena and given 5 min to adapt to baseline illumination (20 µW/cm^2^). After adaption a 15 x 15 mm ROI at the center of the arena was monitored by DAQtimer. Once the larva entered the ROI, full field illumination was extinguished and real-time tracking of the position and orientation of the larva was started. After 3 seconds, two equal intensity light spots (20 µW/cm^2^, 6 mm radius) were positioned 10 mm from the head of the larva at 55° from its current orientation. Light spots were projected onto the base of the arena (AAXA P2 Pico Projector). At the same time, a high-speed camera was triggered to capture the response of the larva to the phototaxis stimuli. Each larva was tested four times with 5 min constant illumination between trials. Trials were excluded from analysis if a phototaxis target was obscured by the environment perimeter or mis-positioned due to sudden movement of the larva, and only larvae with at least 2 trials were analyzed.

#### Multi-day bias persistence tests

(1) For testing individual motor asymmetry over 24 h we first classified individuals as left or right biased at 6 dpf using kinematic routine turn analysis as described above. Classification was determined using the directional preference on the first trial only. Larvae were individually housed and returned to the incubator overnight. We then re-tested larvae at 7 dpf, using the same assay. (2) To determine if motor asymmetry persisted over longer timelines, larvae were tested at 6 dpf using path trajectory analysis over 4 dark trials. Individuals performing same-direction circling for all 4 trials were were then individually housed, and starting at 7 dpf fed daily (AP100 larvae dry food, Zeigler) with the media partially replaced 3 hours after each feeding. No food was given on the day of testing. At 10 dpf larvae were retested using the same assay.

#### Heritability analysis

We selectively raised larvae classified as left or right-biased over two generations. In each generation parents were incrossed and larvae were tested using kinematic analysis (four repeated trials). To enrich for individuals with consistent motor asymmetry, we raised only larvae that (1) had a percentage rightward turn use in the top or bottom quartile of responses and (2) performed all four trials in the same direction. For each generation, left/right classified adults were incrossed as groups and a minimum of 3 independent clutches combined for analysis.

### Calcium imaging

We co-injected 50 ng of *UAS:nls-GCaMP6s* construct (derived from a previously published plasmid *UAS:nls-GCaMP6s-2a-nls-dsRed* (Tabor et al., 2018)) with 80 ng tol1 RNA into one cell stage *y279:Gal4* embryos, and raised in medium containing 200 µM PTU. At 6 dpf larvae were mounted in 2% low melting temp agarose and imaged using a 20x immersion objective on an upright Leica TCS-SP5II microscope. To avoid visual stimulation during GCaMP imaging we used a 2-photon Spectra-Physics MaiTai DeepSee laser tuned to 950 nm and installed a 620 nm LED (All Electronics) on the stage to provide visual stimulation at 80 µW/cm^2^. We performed three trials per larva separated by 3 minutes of constant red light illumination, with each trial consisting of 60 s light, 60 s dark, 60 s light. We captured a single plane through the PT using 16x line averaging of GCaMP fluorescence at approximately 0.96 Hz. After imaging, larvae were recovered from the agarose and placed in fresh E3h media overnight. At 7 dpf we used trajectory analysis of dark-induced circling to determine left/right identity. We analyzed fluorescence intensity changes by manually outlining an ROI around each neuron in a maximum projection of the time series data, and calculated (F_t_ - F_0_)/F_b_ (ΔF/F) where F_0_ was the mean fluorescence intensity during the first baseline period. Neurons with a ΔF/F greater than 3S over three successive timepoints were classified as responders, where S was the standard deviation of ΔF/F during the first baseline period.

### Ablations

#### Genetic ablations

Nitroreductase ablations were performed as previously described (Horstick et al., 2016). To maximize ablation efficiency Gal4 lines were crossed to *UAS:epNTR-tagRFP* and RFP (+) embryos were raised to maturity. Carriers were in-crossed and embryos were sorted using epifluorescence at 2-3 dpf into NTR (+) red fluorescence or NTR (-) groups. Each group was treated with 7.5 mM metronidazole in E3h media from 3 to 5 dpf. Metronidazole media was refreshed after 24 h. At 5 dpf larvae were returned to fresh E3h media and allowed to recover overnight. At 6 dpf genetically ablated larvae were tested and a subset imaged on a confocal microscope to validate ablation efficacy.

#### Laser ablations

We immobilized *y279:Gal4;UAS:Kaede* larvae with tricaine, and performed laser ablation using a Spectra-Physics MaiTai DeepSee laser (tuned to 800 nm, beam power 2 W) on an upright Leica TCS-SP5II microscope with a 20x immersion objective. For ablation we set the ROI to 1-2 neurons per field of view. Efficient ablation was accompanied by the acute and transient development of a bubble. Larvae were then removed from agar and returned to fresh E3h. Controls were anesthetized and mounted for the same duration as ablated larvae. We performed path trajectory analysis during four dark-induced circling trials per larva. For bilaterally ablated larvae, we classified left/right identity using the first trial, then calculated the mean net turn angle for trials 2-4. For unilaterally ablated larvae, where we aimed to see if ablation imposed a pattern of lateralized behavior, we took the mean net turn angle for all four trials. After behavioral analysis we imaged the ablated region to assess ablation efficiency and analyzed only larvae with near complete absence of targeted neurons (~50% of ablated individuals).

### Genetic mapping

To map the *moe^y606^* mutation, we formed mutant and sibling pools, each consisting of 75 embryos that were derived from a single clutch. We Trizol-extracted total mRNA, purified on a Qiagen RNA mini-cleanup column and sequenced samples using single end 100 bp reads using an Illumina HiSeq (100 M reads per sample). We then used RNAmapper in Galaxy (galaxyproject.org) to perform bulk segregant analysis, identifying a critical region on chromosome 9 and a 7.7-fold reduction in expression of *epb41l5* within that interval. We noted that no reads were present in exon 1 or 2 of *epb41l5*; these exons amplified correctly from sibling genomic DNA but not from mutant DNA (exon 1 primers: 5’-TCCACTTTTGGGGATTTACG, 5’-ATTCAATGGCGGAGCAATAC; exon 2 primers: 5’-GGCCATTGACAGTAGTGTGG, 5’-TGACAAGACGCTGAACAAGC) suggesting the presence of a deletion. We then used PCR to assess the presence of genomic DNA in mutants in intervals across a 20 kbp candidate region from between exon 3 of *epb41l5* to the 3’UTR of the neighboring gene *ptpn4a*. This narrowed the candidate region to 7 kb allowing us to design primers to amplify and sequence across the deletion, revealing loss of genomic DNA in mutants between bases chr9:213682 to 218117 (GRCz11, chr9_KZ114909v1_alt). In complementation testing with *moe^b476^*, we recovered the curly-up phenotype in 50/192 (26%) embryos, confirming that this was the causative mutation. Sequencing of the PCR product that spanned the deletion enabled us to design a genotyping protocol to distinguish wildtype and heterozygous larvae using 3 primers (5’-CTACCTGAACAAACTCAATCCAGTC, 5’-AACCATAATAAAATGAGCGTCTCT and 5’-TCATTTTGAAATGCCTGCAA). These primers amplify a single 296 bp band from the wildtype locus, and a 333 bp band from embryos with the deletion.

Previous work established the presence of a large deletion in *moe^b476^* but did not define its precise boundaries, precluding genotyping of heterozygotes (Jensen et al., 2001). To map the *moe^b476^* deletion we used whole-genome shotgun sequencing with 100 bp paired-end reads. After mapping reads to GRCz11 with bowtie2, we focused on a large read-depleted region on chr9 covering *epb41l5*. IGVviewer revealed 5 reads where the paired end reads spanned the read-depleted region. PCR across the putative deletion using primers 5-AAACTGCATAAGTGCCTCACC and 5-GAGACATCGATTCCGCTTTG amplified a ~600 bp band. Sequencing of this band produced the expected genomic sequence at each end and demonstrated that the deletion in *moe^b476^* spans chr9:28835868 to 29370835, completely deleting *epb41l5*, *ptpn4a*, *tmem177* and *pth2r*, and exons from *zgc:91818* and *hs6st3b*. We then used these PCR primers to genotype *moe^b476^* outcrosses because only heterozygous larvae yielded the 600 bp band.

#### Genotyping

After behavioral analysis we genotyped *moe^y606^* and *moe^b476^* larvae as described above. *mib^ta52b^* larvae were genotyped by PCR amplification across the mutation (primers: 5’ GGTGTGTCTGGATCGTCTGAAGAAC; 5’ GATGGATGTGGTAACACTGATGACTC, product size 194 bp). To discriminate wildtype from heterozygous larvae PCR products were digested with the restriction enzyme NlaIII (New England Biolabs) which digests wildtype but not mutant sequence (WT 155 bp; Het 155, 194 bp; Mutant 194 bp). For experiments with otpa mutants, we compared the ‘cousin’ offspring of wildtype (+/+) and mutant (-/-) sibling parents.

### Photoconversion

We characterized projections from *y279-Gal4, UAS:Kaede* expressing rostral PT neurons by focal photoconversion of Kaede from green to red fluorescence in one PT hemisphere. At 2 dpf PT neurons constitute a salient bilateral cluster in the ventral diencephalon. To photoconvert Kaede, larvae were mounted in 2% low melting temperature agarose on an upright Leica TCS SP5 II confocal microscope. We used a 25x immersion objective and set the field of view to comprise 1-2 Kaede expressing PT neurons, imaged with a 488 nm laser. For photoconversion, we performed a single scan of the field of view with 8 frame averaging using a 405 nm laser at 5% power. Conversion of Kaede to the red fluorescent state was confirmed by imaging with a 568 nm laser. After photoconversion embryos were removed from agarose and maintained in E3h until 5 or 6 dpf for imaging the PT and projections to the habenula.

### Neuron Counts

We counted *y279:Gal4, UAS:Kaede* expressing neurons in the rostral PT in larvae imaged at 6 dpf on an upright Leica TCS SP5 II confocal using the 488 nm laser, a 25x immersion objective and 3x zoom. Images were imported into Imaris for counting. After imaging, larvae were extracted from the agar, individually maintained in 6 well plates for 24 h, then tested at 7 dpf to determine left/right identity using path trajectory analysis of dark-induced circling behavior.

### Statistical analysis

Analysis was performed in IDL (Harris), RStudio (Mathworks) and Gnumeric (http://projects.gnome.org/gnumeric/). Data in figures and text are means ± SEM except where otherwise noted. All t-tests were 2-sided. N reported in figure legends. Box plots show median and quartiles with whiskers indicating 10-90%. Normality was determined by the Shapiro-Francia test. Analysis of non-normal data sets was performed using the Mann-Whitney U-test or for one-sample comparisons to a given number, through Monte Carlo testing (see below).

To analyze whether distributions of turn direction deviated from that which would be expected if larvae did not show direction bias, we used Monte Carlo simulations rather than comparison to a binomial distribution because the direction of sequential turns are not independent (Chen and Engert, 2014; Horstick et al., 2017). In each simulation, we used the same number of larvae as in the experimental data (i.e., 89 and 39 respectively for light-off and baseline conditions), and the same number of routine-turns produced on each of the four trials for each larva. As sequential routine-turns show a statistically significant likelihood to be executed in the same direction we simulated routine-turn direction across events on a given trial by randomly selecting a turn-direction for the first event, then then using the previously measured ‘lock index’ to weight a random decision on whether each subsequent turn would be in the same or reverse direction. We used lock indices of 62.8 and 13.2 for light-off and baseline simulations respectively (corresponding to 81.4% and 56.6% chance of executing sequential same-direction routine-turns)(Horstick et al., 2017). The expected histograms of %rightward turns in Figure 1 are derived from 10,000 simulations. In the simulation data, 10% of larvae showed less than 23% right-ward turns or greater than 76% right-ward turns during light-off conditions (i.e., a modal value of 9 of the 89 larvae), whereas in our data, 37 of the 89 larvae exceeded these thresholds. This proportion was greater than all 10,000 simulations, hence we assign a p-value of < 0.0001. In comparison, under baseline conditions, simulations 10% of larvae showed less than 31% rightward turns or greater than 68% right-ward turns during light-off conditions, and in our data 7 of the 39 larvae exceeded these thresholds. However, this proportion was exceeded by 1195 simulations, hence we assign a p-value 0.119. We used a similar procedure to compare the means of larval Match Indices to 0.5, by ranking the difference between the mean of actual match indices in experimental data, with means derived from 100,000 simulations using the same number of larvae and trials per larvae under the null hypothesis assumption that larvae had a 50% (i.e. random) probability of matching the trial 1 direction on each of trials 2-4.

### Data and Software Availability

Further information and requests for image datasets and analysis software should be directed to Harold Burgess or Eric Horstick

