## Supplemental Figures 1-7 for "Molecular and cellular determinants of motor asymmetry in zebrafish"

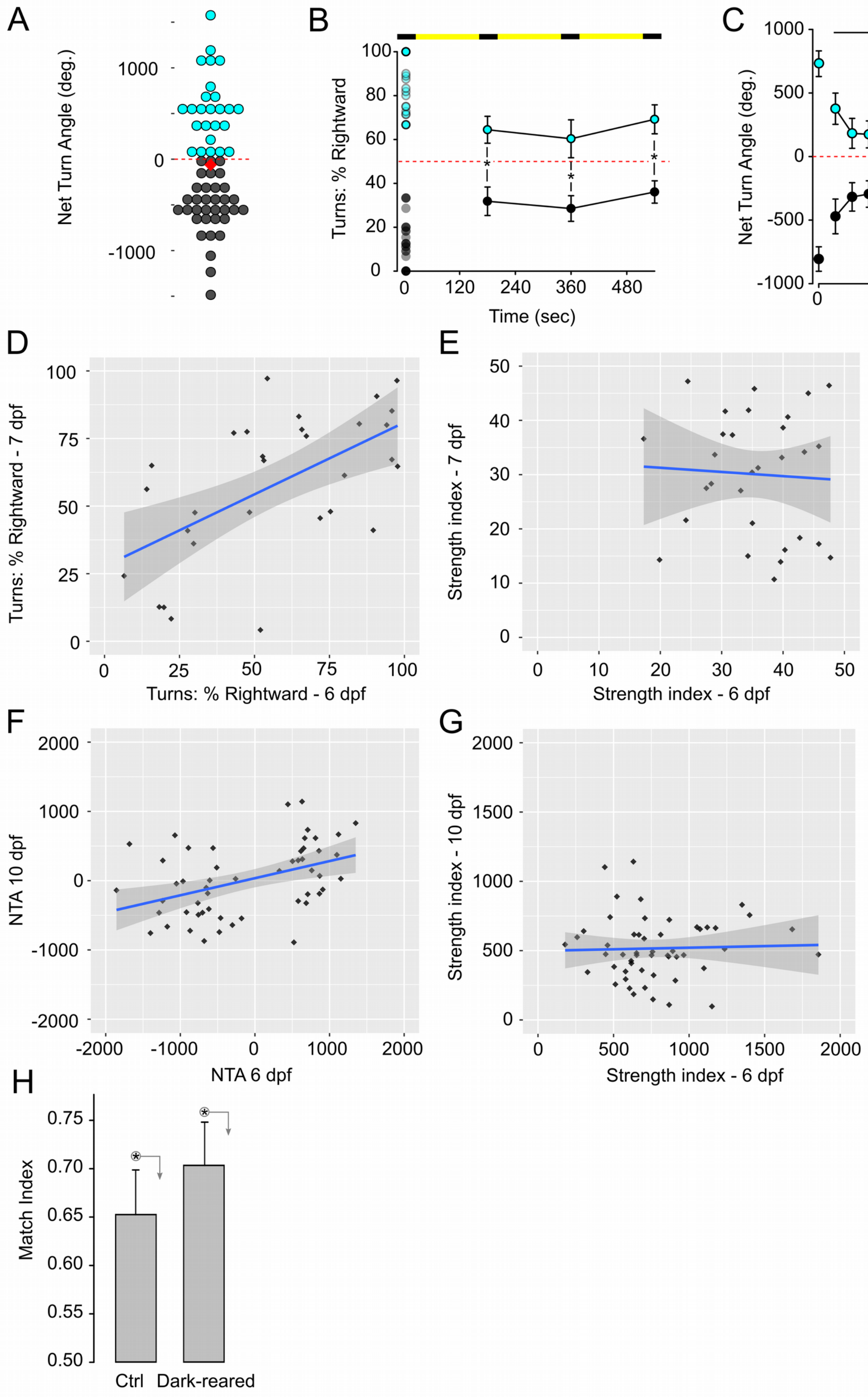
A
Net Turn Angle (deg.)
1000
0
-1000
B
Turns: % Rightward
100
80
60
40
20
0
0
120
240
360
480
Time (sec)
C
Net Turn Angle (deg.)
1000
500
0
-500
-1000
0
20
40
Time (min)
D
Turns: % Rightward - 7 dpf
Turns: % Rightward - 6 dpf
E
Strength index - 7 dpf
Strength index - 6 dpf
F
NTA 10 dpf
NTA 6 dpf
G
Strength index - 10 dpf
Strength index - 6 dpf
H
Match Index
0.75
0.70
0.65
0.60
0.55
0.50
Ctrl
Dark-reared

Figure S1

**Figure S1. Zebrafish larvae exhibit left/right motor bias.**

A. Dotplot of Net Turn Angle during the first 30 s interval after loss of illumination for larvae in Figure 1B. Colors indicate bias (right, cyan, N=25 ; left, grey, N=34). Diamond, population mean.

B. Percentage of routine turns made in a rightward direction for each of four trials. Larvae were classified as right (cyan, N=22) or left (grey, N=24) biased based on the first trial (dotplot at time 0). Larvae with less than 33% of turns in a rightward direction were classified as left-biased, and those with greater than 66% classified as right-biased. The means of the left and right groups are shown for trials 2-4. * $p < 0.05$ between groups.

C. Net turn angle for 16 dark trials for larvae classified based on first trial responses as right (cyan, N=17) or left (grey, N=24) bias. Each dark trial lasted 30 s, with 180 s of illumination between trials. * $p < 0.05$ between groups.

D. Percentage of turns made in a rightward direction (mean of 4 trials) for larvae tested at 6 dpf and again at 7 dpf (N=30).

E. Strength index of routine-turn direction bias for the larvae tested in (D). Index is calculated by taking the absolute difference between the percentage of routine turns executed in a rightward direction and 50% (e.g. 30% rightward turns would give a value of 20 ; 100% would give 50) for each of the four trials per larva, then taking the mean of the four values. This index measures the 'commitment' of a larvae to circle in a given direction on each day.

F. Net turn angles (mean of four trials) for larvae tested at 6 dpf and 10 dpf (N=54).

G. Strength index of direction bias for larvae tested in (F). Index is the mean of the absolute values of the NTA for each of the four trials per larva

H. Match index for control (N=47) and dark-reared (N=41) larvae. * $p < 0.05$, permutation analysis.

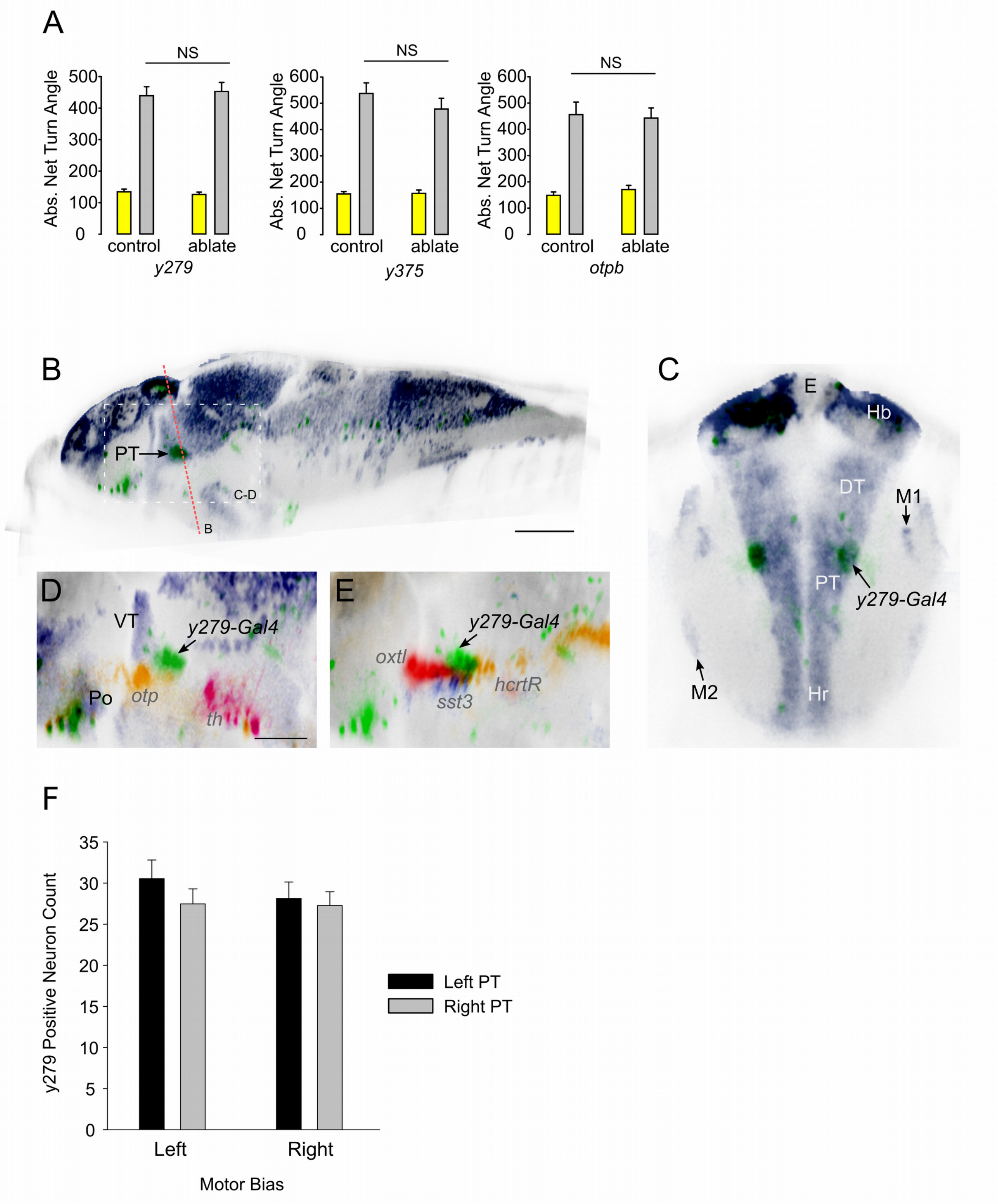
A
NS
Abs. Net Turn Angle
0
100
200
300
400
500
600
control
ablate
y279
y375
otpb
B
PT
C-D
B
C
E
Hb
DT
M1
PT
y279-Gal4
M2
Hr
D
VT
y279-Gal4
Po
otp
th
E
y279-Gal4
oxtl
sst3
hcrtR
F
y279 Positive Neuron Count
0
5
10
15
20
25
30
35
Left
Right
Motor Bias
Left PT
Right PT

Figure S2

**Figure S2.** ***y279-Gal4*** **expressing neurons in the medial diencephalon are part of the posterior tuberculum.**

A. Total turning after chemogenetic ablation for drug treated controls (*y279,* N=54 ; *y375,* N=43 ; *otpbA,* N=35) and following genetic ablation (*y279* N=58 ; *y375* N=37 ; *otpbA* N=46) during paired baseline (yellow) and dark (grey) responses.

B-C. We identified the medial diencephalic cluster of neurons in *y279-Gal4* as within the posterior tuberculum by comparison to the Mueller and Wullimann atlas and Z-Brain (Mueller and Wullimann, 2005; Randlett et al., 2015). In the online version of ZBB (zbbrowser.com) these neurons are located at coordinates 272,307,209 (sagittal, transverse, horizontal planes). ZBB sagittal (B) and transverse (C) sections (sagittal 272 ; transverse 270 with oblique angle of 12 deg) displaying *huC:nls-mCar* (purple) and *y279-Gal4* (green) for comparison to the Mueller and Wullimann atlas Hu-protein antistain in 5 dpf brain panel 31 (page 125). The *y279* neurons are within the central hu$^{+}$ domain, between the M1 and M2 migrated cell groups annotated by Mueller and Wullimann as the ventral posterior tuberculum. We confirm a previous finding that PT neurons project to the habenula, although in the previous report, habenula-projecting PT neurons were assigned to M2 region which is lateral to the *y279* cluster (Hendricks and Jesuthasan, 2007). Scale bar 100 µm.

D-E. Same ZBB sagittal section with (D) *y279-Gal4* (green), *th:Gal4* (pink), *otpB.A:Gal4* (orange) and *gad1b:GFP* (blue). (E) *y279-Gal4* (green), *sst3:Gal4* (blue), *hcrtR:Gal4* (orange) and *oxtl:GFP* (blue). Mid-diencephalic *y279* neurons do not overlap with the GABA transgenic marker gad1b:GFP, and are therefore unlikely to be part of the ventral thalamus (Mueller, 2012). The *y279* cluster is within the PT mask in Z-Brain, although a small section at the dorsal aspect is assigned to the neighboring preoptic area (Randlett et al., 2015). Thus, we assign *y279* neurons to a non-dopaminergic rostral lobe derived from the embryonic ventral PT. Developmentally, the PT originates from the basal plate of prosomeres 1-3 and ventral PT specifically from prosomere 3 (Vernier and Wullimann, 2009). Teleost PT dopaminergic neurons and the mammalian dopaminergic neurons of the substantia nigra/ventral tegmental area are likely homologous. Mammalian non-dopaminergic derivatives of basal prosomeres are less clear but the basal plate of prosomere 3 gives rise to the Fields of Forel (Puelles and Rubenstein, 2003). Scale bar 50 µm. DT, thalamus. E, epiphysis. Hb, habenula. Hr, rostral hypothalamus. M1, migrated pretectal area. M2, migrated posterior tubercular area. Po, preoptic region. PT, posterior tuberculum. VT, ventral thalamus.

F. *y279-Gal4* expressing rostral PT neuron counts (left hemisphere black bar ; right hemisphere grey bar) in left (N=13) and right (N=15) biased larvae.

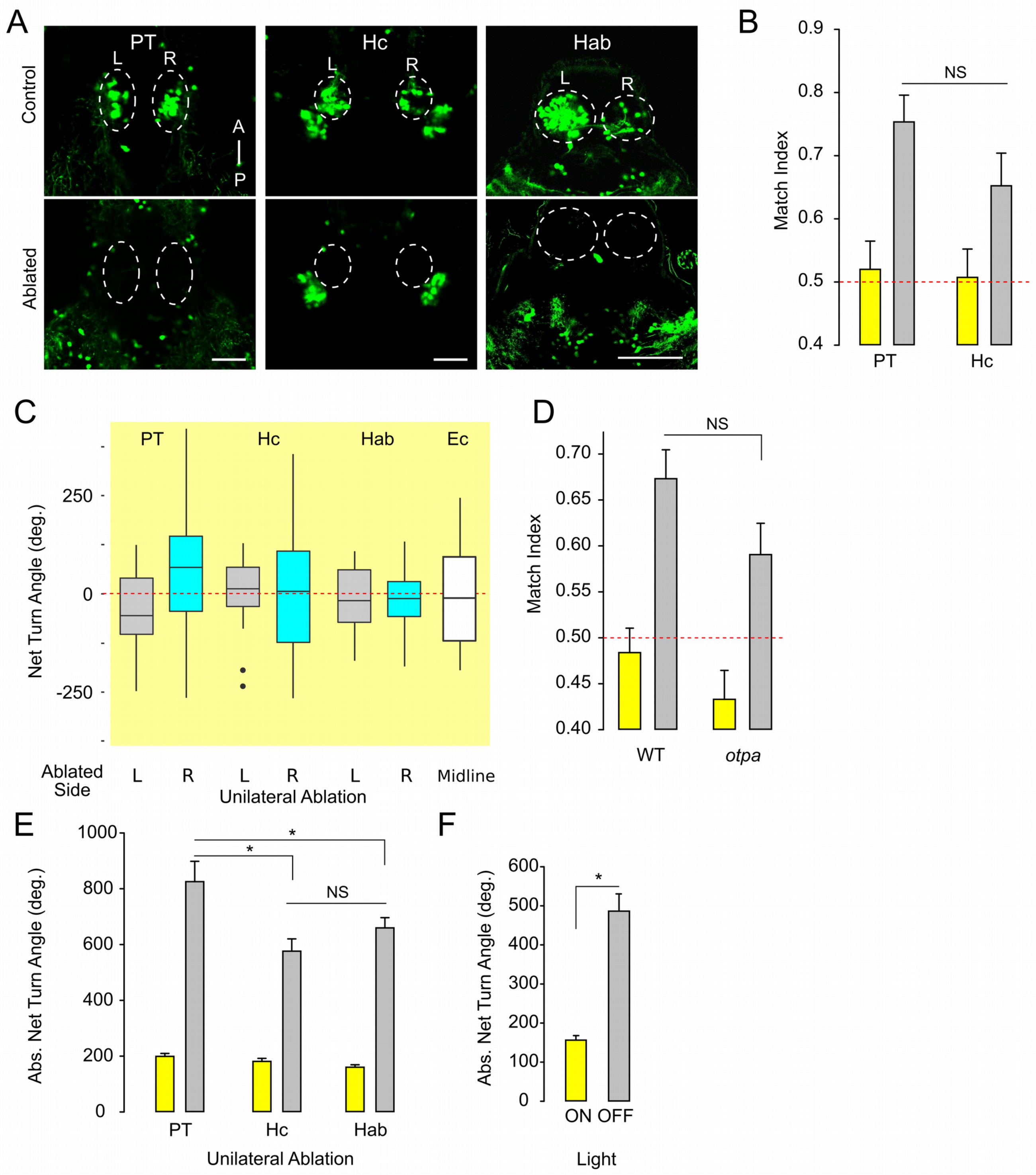
A
PT
Hc
Hab
Control
Ablated
L
R
A
P
B
Match Index
0.9
0.8
0.7
0.6
0.5
0.4
NS
PT
Hc
C
PT
Hc
Hab
Ec
Net Turn Angle (deg.)
250
0
-250
Ablated Side
L
R
L
R
L
R
Midline
Unilateral Ablation
D
Match Index
0.70
0.65
0.60
0.55
0.50
0.45
0.40
NS
WT
otpa
E
Abs. Net Turn Angle (deg.)
1000
800
600
400
200
0
*
*
NS
PT
Hc
Hab
Unilateral Ablation
F
Abs. Net Turn Angle (deg.)
600
500
400
300
200
100
0
*
ON OFF
Light

Figure S3

**Figure S3. Behavioral analysis after posterior tuberculum and habenula ablations.**

A. Confocal projection of *y279* expressing posterior tuberculum (PT), caudal hypothalamus (Hc) and habenula (Hab) neurons in representative control (top) and laser ablated (bottom) larvae. Scale bar 50 µm.

B. Match index after unilateral ablations of *y279* posterior tuberculum (PT, N=50) or caudal hypothalamus neurons (Hc, N=46) during baseline illumination (yellow) or dark-induced circling (grey). Left and right hemisphere ablations are combined in this analysis.

C. Net turn angle (mean of trials 1-4) under baseline illumination after unilateral ablation of left (grey, PT N=27 ; Hc N=24 ; Hab N=33) or right (cyan, PT N=23 ; Hc N=22 ; Hab N=28) hemisphere *y279* PT or Hc neurons. Baseline net turn angle after laser section of the epithalamic commissure (Ec) (white, N=24).

D. Match index for cousin wildtype (N=103) and *otpa* mutant larvae (N=92). Dotted red line denotes random motor bias.

E. Total turning after unilateral laser ablations of *y279* posterior tuberculum (PT, N=50), caudal hypothalamus (Hc, N=46) or habenula (Hab, N=61) neurons under baseline illumination (yellow) and during dark trials (grey). Left and right hemisphere ablations are combined in this analysis. * $p < 0.05$.

F. Total turning behavior following Ec section for larvae in Figure 3G, S3C. t-test * $p < 0.05$.

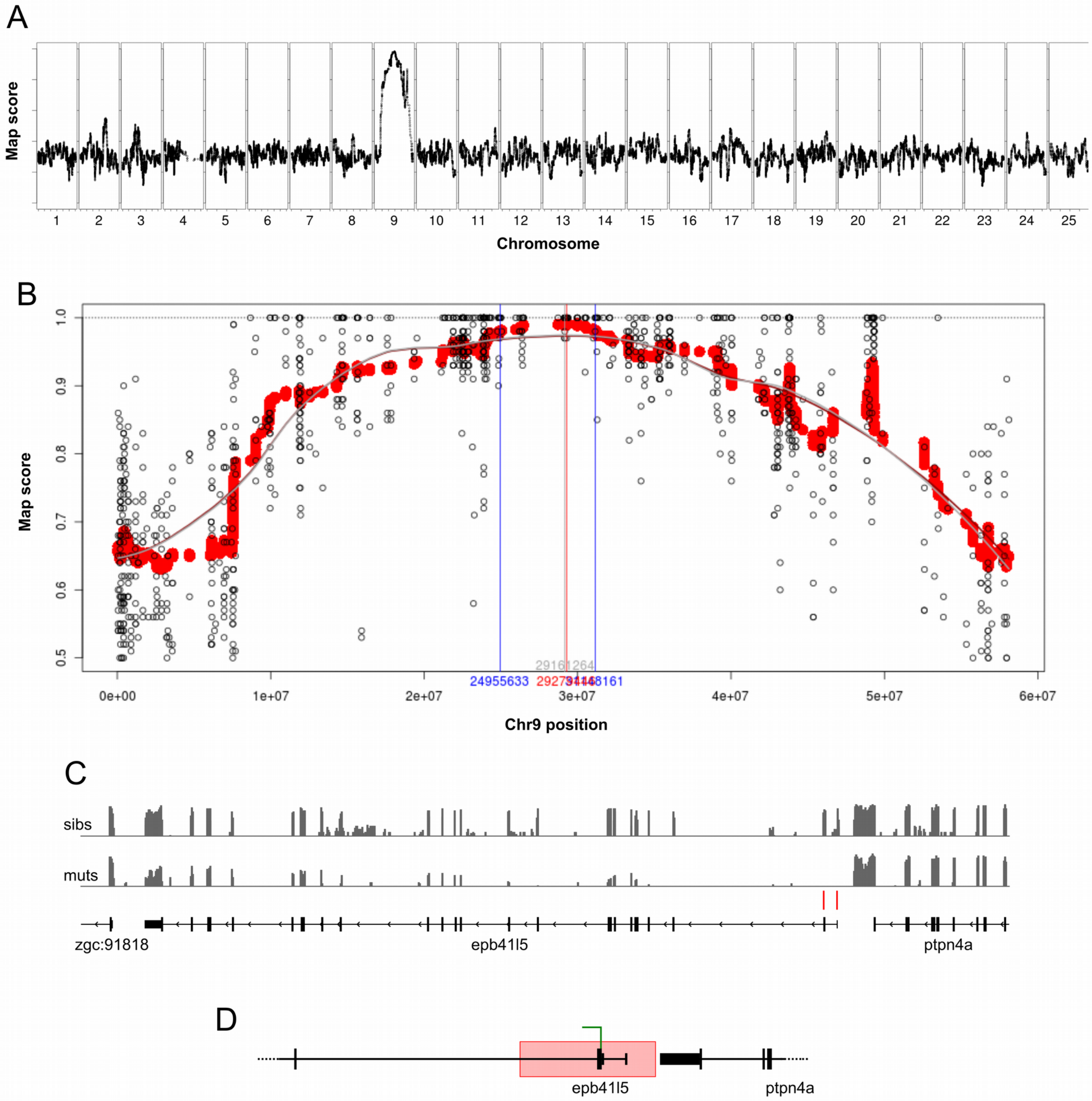
A
Map score
Chromosome
B
Map score
1.0
0.9
0.8
0.7
0.6
0.5
24955633
0e+00
1e+07
2e+07
3e+07
4e+07
5e+07
6e+07
Chr9 position
C
sibs
muts
zgc:91818
epb41l5
ptpn4a
D
epb41l5
ptpn4a

Figure S4

**Figure S4. Mapping of the *moe*$^{y606}$ mutation.**

A. Whole genome Galaxy RNAmapper pipeline map scores, comparing pooled mutants with the curly-up phenotype and siblings

B. Map scores for chromosome 9.

C. Read counts (IGVtools count function) from bulk RNAseq for curly-up mutants and siblings. Arrows indicate a complete lack of reads for exons 1 and 2 of *epb41l5* (one read mapped to exon 3).

D. Schematic of the deletion in *moe*$^{y606}$ which eliminates the first two exons of *epb41l5*. The deletion removes bases 213682-218117 from chr9_KZ114909v1_alt (an alternative build for part of chromosome 9) and inserts 53 low complexity bases that are not found in this region.

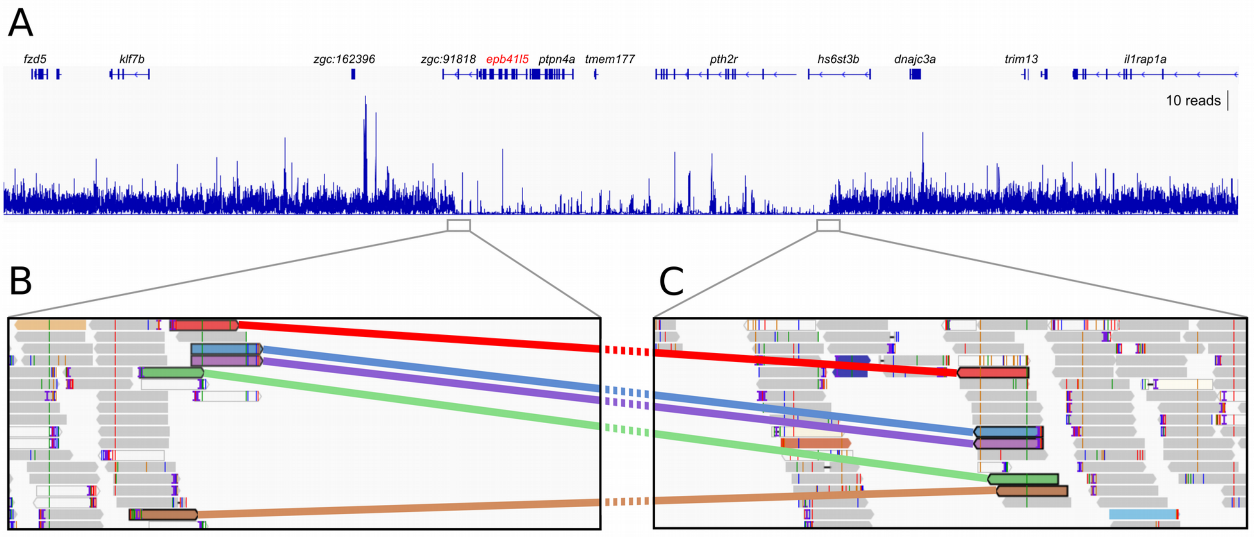
A
fzd5
klf7b
zgc:162396
zgc:91818
epb41l5
ptpn4a
tmem177
pth2r
hs6st3b
dnajc3a
trim13
il1rap1a
10 reads
B
C

Figure S5

**Figure S5. Mapping of the *moe*$^{b476}$ deletion.**

A. RNAseq read mapping from *moe*$^{b476}$ mutant larvae, in the region chr9:28,190,541 to 29,960,696. Grey boxes indicate the areas highlighted in (B).

B-C. RNAseq reads mapped to the left side of the deletion (B, chr9:28,835,314 to 28,836,377) and right side of the deletion (C, chr9:29,370,348 to 29,371,411). The 6 colored reads indicate reads where one of the paired ends mapped to one side of the deletion, and the other paired end mapped to the other side of the deletion.

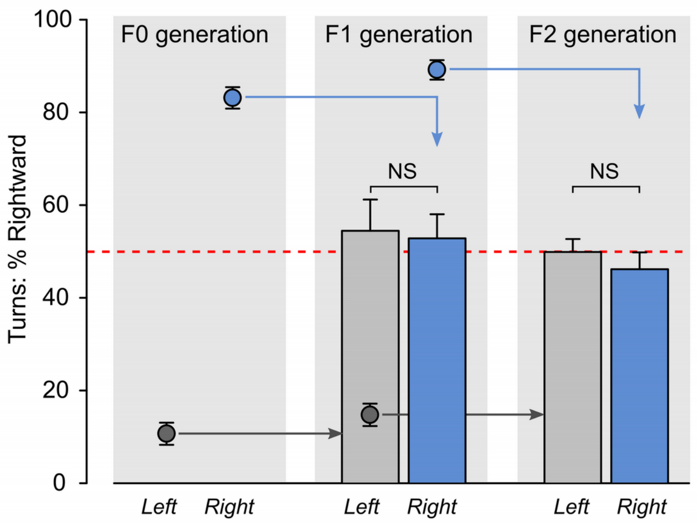

100
80
60
40
20
0
Turns: % Rightward
F0 generation
F1 generation
F2 generation
NS
NS
Left
Right
Left
Right
Left
Right

Figure S6

**Figure S6. Left-right bias is not heritable.**

Mean percent rightward turns for groups of larvae used to assess heritability of left/right motor asymmetry. F0 generation: circles indicate mean values of larvae selected to raise to become parents of the F1 generation (left, N=10 ; right, N=8). F1 generation: bars indicate the population means for F1 larvae derived from left-biased (grey, N=31) or right-biased (blue, N=37) F0 adults. Circles indicate mean values of larvae raised as parents of the F2 generation (left, N=7 ; right, N=9). F2 generation: bars indicate populations means for F2 larvae (left, N=86 ; right, N=69).

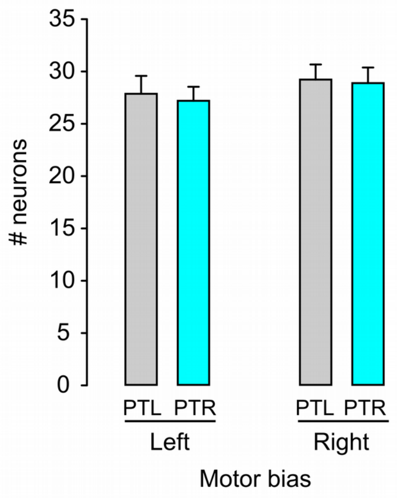
35
30
25
20
15
10
5
0
### neurons
PTL
PTR
PTL
PTR
Left
Right
Motor bias

Figure S7

**Figure S7. Neuron counts in left/right posterior tuberculum in heterozygous *moe*$^{y606}$ larvae.**

Number of *y279-Gal4, UAS:Kaede* expressing neurons in left (N=16) and right (N=10) biased *moe*$^{y606/+}$ larvae in the left PT (grey) and right PT (cyan).
